# The SEQC2 Epigenomics Quality Control (EpiQC) Study: Comprehensive Characterization of Epigenetic Methods, Reproducibility, and Quantification

**DOI:** 10.1101/2020.12.14.421529

**Authors:** Jonathan Foox, Jessica Nordlund, Claudia Lalancette, Ting Gong, Michelle Lacey, Samantha Lent, Bradley W. Langhorst, V K Chaithanya Ponnaluri, Louise Williams, Karthik Ramaswamy Padmanabhan, Raymond Cavalcante, Anders Lundmark, Daniel Butler, Chris Mozsary, Justin Gurvitch, John M. Greally, Masako Suzuki, Mark Menor, Masaki Nasu, Alicia Alonso, Caroline Sheridan, Andreas Scherer, Stephen Bruinsma, Gosia Golda, Agata Muszynska, Paweł P. Łabaj, Matthew A. Campbell, Frank Wos, Amanda Raine, Ulrika Liljedahl, Tomas Axelsson, Charles Wang, Zhong Chen, Zhaowei Yang, Jing Li, Xiaopeng Yang, Hongwei Wang, Ari Melnick, Shang Guo, Alexander Blume, Vedran Franke, Inmaculada Ibanez de Caceres, Carlos Rodriguez-Antolin, Rocio Rosas, Justin Wade Davis, Jennifer Ishii, Dalila B. Megherbi, Wenming Xiao, Will Liao, Joshua Xu, Huixiao Hong, Baitang Ning, Weida Tong, Altuna Akalin, Yunliang Wang, Youping Deng, Christopher E. Mason

## Abstract

Cytosine modifications in DNA such as 5-methylcytosine (5mC) underlie a broad range of developmental processes, maintain cellular lineage specification, and can define or stratify cancer and other diseases. However, the wide variety of approaches available to interrogate these modifications has created a need for harmonized materials, methods, and rigorous benchmarking to improve genome-wide methylome sequencing applications in clinical and basic research. Here, we present a multi-platform assessment and a global resource for epigenetics research from the FDA’s Epigenomics Quality Control (EpiQC) Group. The study design leverages seven human cell lines that are designated as reference materials and publicly available from the National Institute of Standards and Technology (NIST) and Genome in a Bottle (GIAB) consortium. These samples were subject to a variety of genome-wide methylation interrogation approaches across six independent laboratories, with a primary focus was on 5-methylcytosine modifications. Each sample was processed in two or more technical replicates by three whole-genome bisulfite sequencing (WGBS) protocols (TruSeq DNA methylation, Accel-NGS MethylSeq, and SPLAT), oxidative bisulfite sequencing (TrueMethyl), one enzymatic deamination method (EMseq), targeted methylation sequencing (Illumina Methyl Capture EPIC), and single-molecule long-read nanopore sequencing from Oxford Nanopore Technologies. After rigorous quality assessment and comparison to Illumina EPIC methylation microarrays and testing on a range of algorithms (Bismark, BitmapperBS, BWAMeth, and GemBS), we found overall high concordance between assays (R=0.87-R0.93), differences in efficency of read mapping and CpG capture and coverage, and platform performance. The data provided herein can guide continued used of these reference materials in epigenomics assays, as well as provide best practices for epigenomics research and experimental design in future studies.

## Introduction

DNA methylation plays a key role in the regulation of gene expression [1], disease onset [2], cellular development [1], age progression [3], and transposable element activity [4]. Whole Genome Bisulfite Sequencing (WGBS) is increasingly used for fundamental and clinical research of CpG methylation. Numerous validated protocols and commercially available kits are available for WGBS library preparation ([5], [6], [7]). Other assays to interrogate the epigenome include oxidative bisulfite sequencing [8], enzymatic deamination [9], and targeted approaches ([10], [11]), further accelerating the breadth and rate of discovery in genome-wide DNA methylation studies.

As the field of epigenomics continues to advance, there is a need to establish definitive standards and benchmarks repesentative of the methylome. The Genome in a Bottle (GIAB) Consortium has recently established seven human cell lines as reference material to enable genomics benchmarking and discovery [12]. Recent work has characterized the genomes of these cell lines (e.g. germline structural variant detection in [13]), but not yet at the epigenome level. Here, the FDA’s Epigenomics Quality Control (EpiQC) Group presents epigenomic sequence data across all seven GIAB reference cell lines, as well as a comparative analysis of targeted and genome-wide methylation protocols, to serve as a comprehensive resource for epigenetics research. We build on top of previous work done to compare the performance and biases of WGBS library kits (e.g. [6, 14, 15]) by evaluating both commonly used and newly available epigenomic library preparation kits across a broad set of samples that are used increasingly for benchmarking. We report the relative performance of each kit, as measured by mapping efficiencies, CpG coverage, and methylation estimates, as well as characterizing the reproducibility and challenges of methylation estimation across the genome. We further sequenced these cell lines using long read technology on an Oxford Nanopore PromethION and compare its ability to characterize the epigenome alongside more common chemical/enzymatic conversion kits and short read sequencing. We also generated microarray data for these cell lines and provide guidelines for normalization of beta values, site filtration, and comparison to epigenetic sequence data. This reference dataset can act as a benchmarking resource and a reference point for future studies as epigenetics research becomes more widespread within the field of genomics.

## Results

### Study Design and Sequencing Outputs

We generated epigenomic data for seven well-characterized human cell lines (HG001-HG007) that have recently been designated as reference materials for genomic benchmarking by the Genome in a Bottle (GIAB) Consortium [16]. These cell lines include NA12878 (HG001) from the CEPH Utah Reference Collection, as well as two family trios from the Personal Genome Project, one of Ashkenazi Jewish ancestry (HG002-4) and one of Han Chinese ancestry (HG005-7).

Libraries for whole epigenome sequencing were prepared using a variety of common bisulfite and enzymatic conversion kits, including NEBNext Enzymatic Methyl-Seq (referred to here as EMSeq), Swift Biosciences Accel-NGS Methyl-Seq (MethylSeq), SPlinted Ligation Adapter Tagging (SPLAT), NuGEN TrueMethyl oxBS-Seq (TrueMethyl), and Illumina TruSeq DNA Methylation (TruSeq). Cell line genomic DNA was acquired from Coriell, and one aliquot of each genome was extracted and distributed to six independent laboratories, each utilizing one library preparation method (Table 1).

**Table 1.**
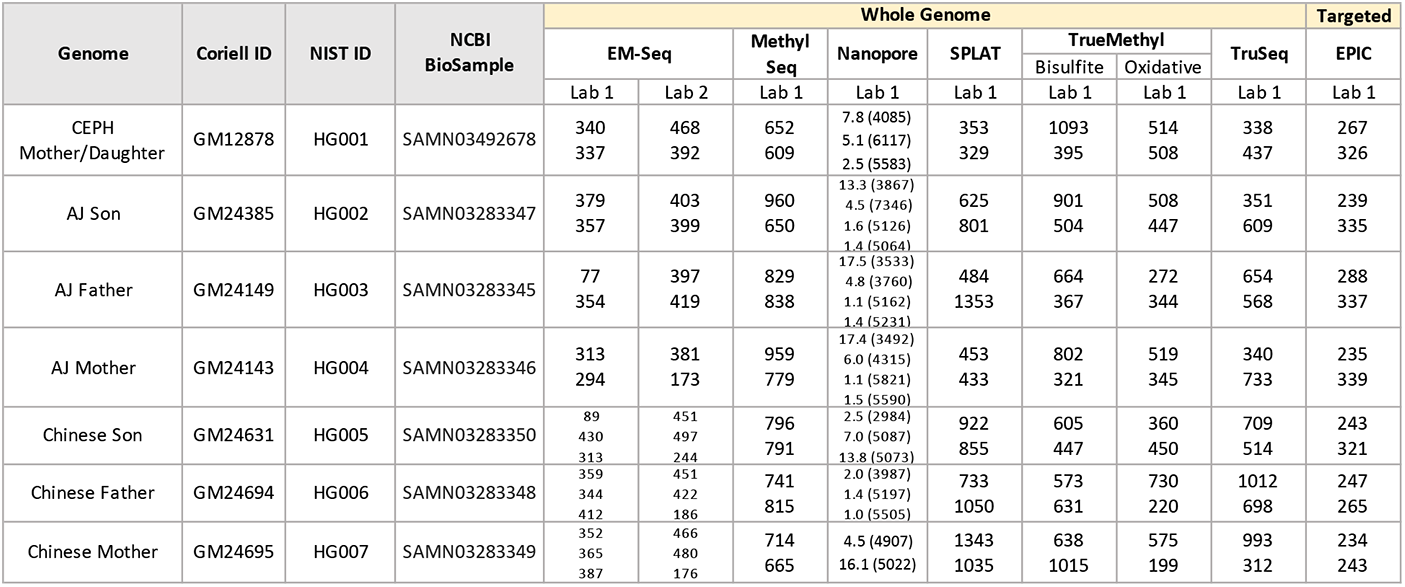
Sequencing across all genomes analyzed in this study, including genomic and targeted assays. Numbers within each genome/assay cell indicate millions of paired-end 150bp reads sequenced, with the exception of PromethION, which indicates millions of reads and mean read length in parentheses. Each number represents one replicate sequenced for that genome/assay.

Each site prepared two technical replicates per cell line for their respective epigenetic assay. In the case of EMSeq, libraries were prepared at two sites, designated as Lab 1 and Lab 2. All other sites were designated as Lab 1 for their library type. In the case of TrueMethyl, pairs of replicates were made using a bisulfite-only treatment (BS) and an oxidative-bisulfite treatment (OX). All libraries were pooled into equamolar concentrations and sequenced in multiplex at one site (see methods), resulting in a range of 500M to 3.5B paired-end reads per replicate. The range of sequencing depth per replicated resulted from an imbalance in library pooling, as well as differences in shearing condition and size selection per library type (see methods).

In addition to short read sequencing of epigenetic libraries, Oxford Nanopore R9.4.1 PromethION flow cells (referred to here as Nanopore) were run to generate long read sequence data for each genome, each ranging from 75B to 250B bases.

### Data Quality Control

We performed quality control of all sequence data generated within this study using FASTQC [17] (see Supplementary Data 1 for quality reports for every sample). As a measure of the success of the bisulfite or enzymatic convesion step of each library preparation, we estimated the cytosine conversion rate across CpG and non-CpG contexts (Figure S1a). CpG methylation levels fell in the expected 45%-65% range across all libraries (Methyl Capture EPIC, as an exception, showed lower rates, a reflection of targeting less methylated regions such as promoters and enhancers). Conversion of cytosines in non-CpG contexts was near zero as expected for all libraries, though CHG and CHH context conversion was somewhat eelevated for TruSeq libraries (Figure S1a) (see below for mapping and methylation calling that enabled these estimates).

Depending on library preparation, different libraries had different completely unmethylated (lambda) or completely methylated (pUC19 plasmid) spiked-in controls (see methods). Methylation levels of these controls were very nearly 0% or 100% respectively across all libraries (Figure S1b), further reflecting the quality of the data.

### Mapping Efficiencies Per Epigenomic Library Type

Following quality control, we examined the performance of reference-based read alignment and methylation estimation for samples of each library type. Our pipeline of choice was bwa-meth (a common methylation-aware, reference-based read aligner) followed by MethylDackel for methylation extraction, which was chosen for its high mapping efficiency, greatest mean depth of coverage per CpG, and speed (for a comparison of alignment and methylation calling pipelines, see the supplementary results, as well as Figure S2 and Figure S3). Each epigenomic assay had a distinct profile of mapping outcomes (Figure 1a). MethylSeq had the highest primary mapping rate and lowest secondary/unmapped rate. While EMSeq (Lab 1) and SPLAT had comparable primary mapping rates to MethylSeq, SPLAT had the highest fraction of unmapped reads. TrueMethyl had the highest rate of multi-mapped reads, while TruSeq returned the highest rate of PCR duplicate reads.

**Figure 1:**
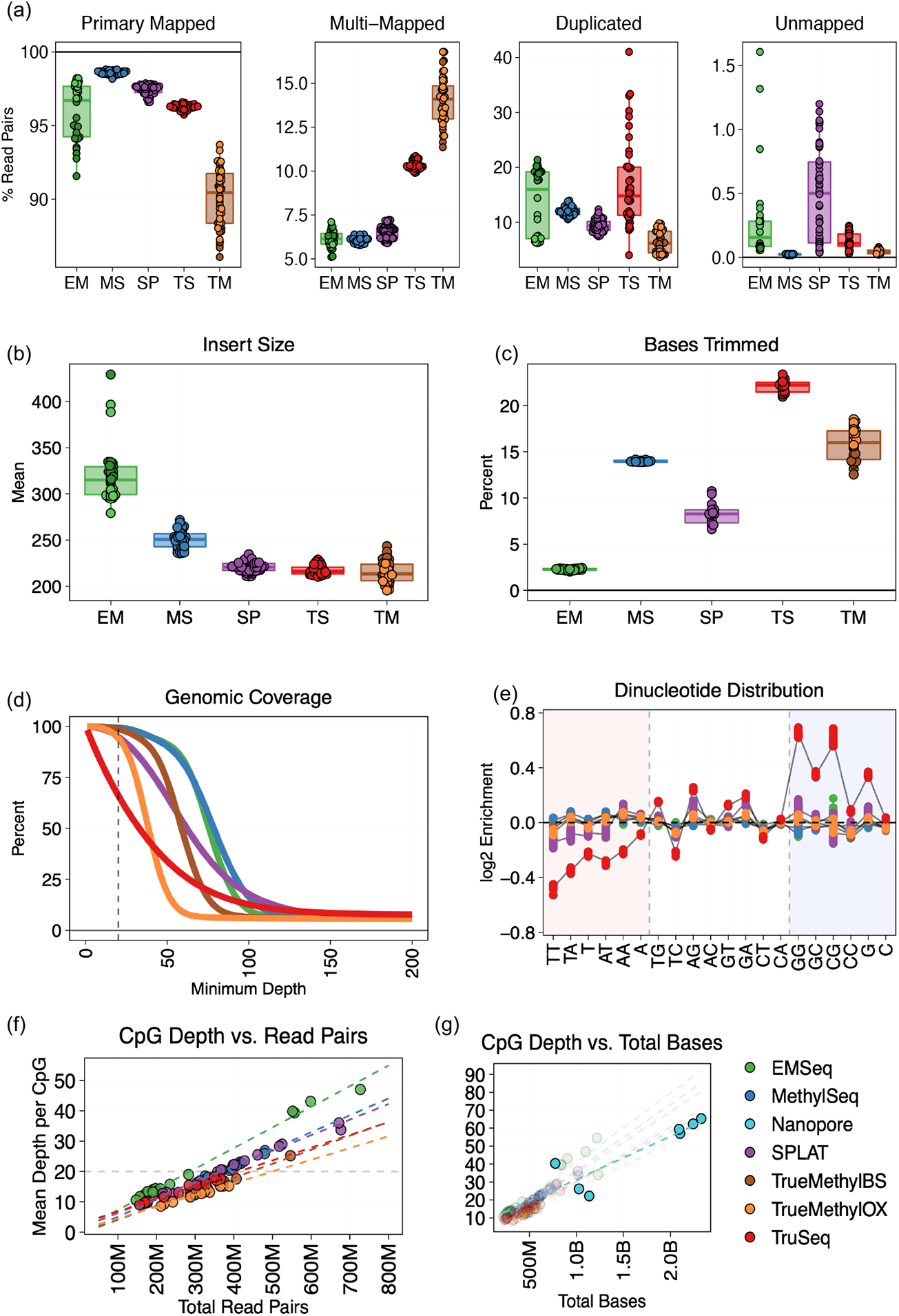
Sequencing and alignment metrics of whole methylome libraries, including all replicates across all cell lines. EM=EMSeq; MS=MethylSeq; SP=SPLAT; TS=TruSeq; TM=TrueMethyl. (a) Distribution of reference-based read alignment outcomes, including primary mapped reads (both mates mapped in correct orientation within a certain distance), multi-mapped reads (read pairs containing secondary or supplementary alignments), reads marked as PCR or optical duplicates, and unmapped reads. Ambiguous and duplicate reads can be a subset of properly aligned reads. (b) Median insert size distributions derived from distance between aligned paired end reads. (c) Percentage of bases trimmed per replicate, either due to low base quality, adapter content, or dovetailing reads. (d) Cumulative genomic coverage plot, averaged across cell line per assay. Coverage is cut off at 200x in this plot, but extends beyond for all assays. (e) Nucleotide bias plot showing the log2 enrichment of covered versus expected mono- and di-nucleotides. (f) The relationship between the number of read pairs sequenced per assay and the mean depth of coverage per CpG dinucleotide, showing sequencing depth required to achieve a certain level of coverage. 20x CpG coverage is shown as the dotted line. (g) Same as (f), but plotted using total bases sequenced, to include Oxford Nanopore sequencing, which produces variable read lengths.

As a measure of protocol efficiency, we estimated the total cytosine conversion in CpG contexts and found that each whole-methylome approach converted 45-65% of CpGs. As an estimate of conversion efficiency, we also characterized methylation in CHG and CHH contexts and found all libraries to be close to the expected 0% range (nearing 100% conversion efficiency), except for TruSeq which neared 2% in CHG contexts and 1% in CHH contexts, and MethylSeq which approached 0.75% in CHH contexts (Figure S1).

Each assay had a specific, tight profile of insert size distributions (Figure 1b). There was a strong relationship within each assay between the estimated insert size and the percentage of total bases that were trimmed prior to alignment (this included triming adapter content, low quality bases, and dovetailing bases between mates of a pair of reads). Libraries with insert sizes below 275bp had anywhere from 5-25% of total bases trimmed, while EMSeq libraries with >275bp insert sizes needed very few bases trimmed other than adapter content (Figure 1c). This particular pattern was seen due to the 150×150 chemistry used for sequencing, and the threshold for fragment size may be lower with shorter read sequencing.

Imbalanced base trimming and unequal distribution of reads per replicate (see above) resulted in divergent genome coverage per assay (Figure 1d). Generally, a minimum of 20X coverage is considered sufficiently deep to characterize a genomic region, and EMSeq and MethylSeq had the highest percentage of the genome covered at 20X. This was followed by SPLAT, the oxidative and bisulfite replicates of TrueMethyl, and lastly the TruSeq libraries, which had the lowest percentage of the genome covered at lower depths, but a long tail of high-coverage sites. TruSeq libraries also showed a high degree of dinucleotide bias favoring GC-rich regions compared to other libraries (Figure 1e), owing to the GC-biased random hexamer ligation step in its library preparation, as well as exposing samples to sodium bisulfite prior to DNA shearing.

Reads from whole methylome libraries were passed through an alignment and methylation calling pipeline (see above). Reads were filtered from the methylation calling process if they did not map to the reference genome, if they were marked as a non-primary alignment (secondary/supplementary/duplicate reads), or if they were assigned a mapping quality score below MQ10. The fractions of reads that were filtered along the alignment pipeline (Figure S4) were highly assay-specific. At the end of this process, EMSeq libraries retained the highest percentage of reads for methylation calling (maximum 86%), followed by SPLAT (83%), MethylSeq (81%), TrueMethyl (80%), and finally TruSeq (77%). EMSeq also showed laboratory specificity, with lower rates of useable bases in libraries prepared using shorter fragment sizes (mean of 86% in Lab 1 versus 73% in Lab 2) (see methods). We observed no notable differences in read filtration rates between TrueMethyl libraries treated with potassium perruthenate (KRuO4) oxidation and those only exposed to sodium bisulfite. The average total percentage of useable bases is summarized per assay for HG002 in Table 2, and more detailed statistics for all cell lines are shown in Supplementary Table 2.

**Table 2.** Summary statistics of mapping and library efficiency per WGBS protocol. Percent CpG capture calculated with call sets normalized to 20x coverage.

|  | EMSeq<br>Lab 1 | EMSeq<br>Lab 2 | Methyl<br>Seq | SPLAT | TrueMethyl<br>(BS) | TrueMethyl<br>(OX) | TruSeq |
| --- | --- | --- | --- | --- | --- | --- | --- |
| Insert Size (bp) | 299 | 327 | 250 | 221 | 224 | 207 | 215 |
| Mapping Rate (%) | 97 | 93 | 98 | 97 | 85 | 86 | 95 |
| Duplicate Rate (%) | 9 | 25 | 12 | 8 | 20 | 20 | 21 |
| Dinucleotide Bias<br>Score | 3 | 1 | 4 | 10 | 4 | 4 | 27 |
| **Useable Bases (%)** | **90** | **77** | **74** | **81** | **70** | **67** | **60** |
| Reads to reach 20x<br>CpG coverage (M) | 275 | 303 | 366 | 369 | 446 | 496 | 692 |
| Mean CpG Depth<br>per replicate | 13, 13 | 13, 13 | 27, 17 | 17, 22 | 20, 15 | 15, 13 | 10, 15 |
| % Genome-wide<br>CpGs ≥ 1x cov | 100 | 100 | 100 | 100 | 100 | 100 | 100 |
| % Genome-wide<br>CpGs ≥ 10x cov | 94 | 92 | 91 | 89 | 91 | 90 | 74 |

We next calculated for each library type the relationship between raw total number of read pairs sequenced versus the mean depth of coverage achieved per CpG (Figure 1f). We found that the rates were highly assay-specific, as seen above. Overall, in order to achieve a target mean depth of 20X per CpG, EM-Seq required the fewest reads (275-300M read pairs), folowed by MethylSeq (366M) and SPLAT (369M), then TruSeq (461M), and then TrueMethyl (692M), as noted in Table 2. In order to compare short read data to variably-lengthed long read data from Oxford Nanopore, we calculated the same relationship using total bases sequenced (Figure 1g). We found that Nanopore sequencing covered CpGs and called methylation at a similar rate per nucleotide as did any short read library type.

### CpG Coverage and Downsampling

We analyzed the distribution of CpG coverage across the genome per assay. In order to control for the effect of uneven sequencing depth, we first downsampled the methylation call sets for every replicate to a given mean coverage value. Downsampling can be done by either filtering the number of reads in an alignment (BAM files), or by randomly removing a fraction of observed cytosines and observed thymines per CpG within methylation call sets (bedGraph files). Because downsampling at the alignment level can be slow and demanding in terms of disk space and compute time, we set out to evaluate if the signal from downsampling cytosines within bedGraph files recapitulated downsampling aligned reads within BAM files. The two approaches yielded similar results in number of CpG sites detected, distribution of read counts, and methylation calls. bedGraph downsampling had the added benefit that the targeted average CpG coverage was more accurately estimated than when downsampling BAMs (Figure S5).

We proceeded with methylation call sets that were normalized to a mean of 20x coverage per site. Unless otherwise noted, these call sets comprised merged replicates per library type, and merged calls on positive and negative strands (i.e. reporting methylaton at the dinucleotide level rather than individual cytosines). The mean coverage per library shifted as expected, indicating the success of the downsampling approach (Figure S6a). Notably, the methylation percentage distribution also shifted, with the bimodal peaks at 0% and 100% becoming more pronounced, and putatively hemimethylated regions dropping out as a function of fewer observations per site resulting in lowered sensitivity (Figure S6b). We observed that downsampling below 20x exaggerated this effect. Downsampling also produced an assay-specific pattern of site dropout (Figure S7). Although the overwhelming number of sites are covered by all assays, we observed the highest CpG dropout in TruSeq, followed by SPLAT, then MethylSeq, then TrueMethyl, then EMSeq, both when accounting for any coverage at all (>=1x) or coverage of >=50% of the overall mean value.

Even after normalizing for mean CpG coverage, we observed a range of assay-specific empirical cumulative distributions (Figure 2a). In particular, TruSeq produced left and right tails of very low and very high coverage. We see this has an effect on reproducibility between replicates of the same assay (Figure 2b), where, compared to an expected distrubtion of cross-replicate concordance, TruSeq showed the highest variation, followed by TrueMethyl, while SPLAT, MethylSeq, and EMSeq were more reproducible than expected. Intra-assay coverage reproducibility was relatively consistent above 20X coverage (r>0.98 for all assays), but broke down below 10X (r<=0.95 for all assays). We therefore recommend 20X as a minimum CpG dinucleotide coverage value (Figure S9).

**Figure 2:**
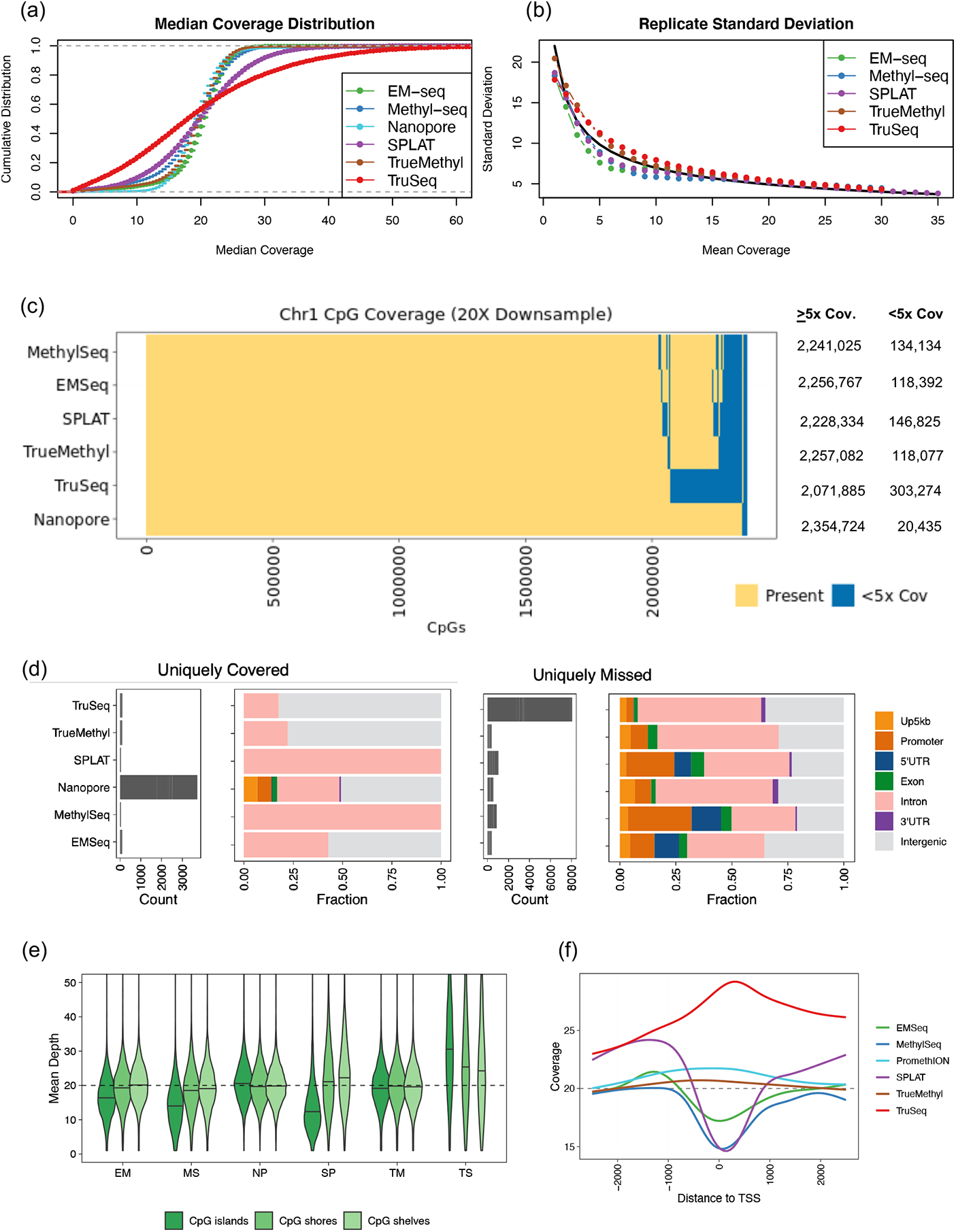
Coverage of CpGs across the genome. All samples visualized here were downsampled to 20X mean coverage per CpG. (a) Empirical cumulative distribution functions for median coverage, averaged across samples for HG002-HG007. (b) Standard deviation between replicate beta values for HG002 as a function of average coverage. The expected curve (computed based on the assumption that replicate beta values are independent and identically distributed estimates of a common proportion p) is added as a solid black curve. (c) Intersection of CpG coverage (min 5x) across Chromosome 1. Exact values of CpGs covered per assay are shown on the right. (d) Count and genomic annotation for CpGs uniquely covered by an assay (left) and uniquely not covered by an assay (right). Up5kb = 5kb upstream distance from promoter region; Promoter = within 1kb upstream of transcript start site. (e) Distribution of coverage in CpG shelves, shores, and islands. EM=EMSeq; MS=MethylSeq; SP=SPLAT; TS=TruSeq; TM=TrueMethyl. (f) Mean coverage curves around transcript start sites (TSS).

We restricted further analyses to Chromosome 1, which represents a significant portion of the genome (10%), contains all difficult regions (such as tandem duplications and satellites), and is computationally much more tractable than a genome-wide analysis. When aligning CpGs covered in the 20X downsampled libraries, we found that the majority of CpGs (>99%) were covered by all assays, with some assay-specific dropout (Figure 2c). Nanopore sequencing was able to cover the highest number of CpGs not covered by other assays, and TruSeq missed the highest number of CpGs covered by other assays (Figure 2d). Among the regions covered uniquely by Nanopore sequencing, about 20% were meaningful for epignetic regulation (promoter, TSS, or exonic sites), while the few CpGs uniquely captured by other assays were intronic or intergenic (Figure 2d). Despite the small number of differences of CpG coverage observed between assays, the total genomic annotation of sites covered was highly consistent (Figure S8).

We also examined the coverage of CpG islands, shelves, and shores (Figure 2e). Nanopore returned the most even coverage across these annotations, while TruSeq showed elevated coverage relative to its overall mean in these GC-rich regions. EMSeq, MethylSeq, and SPLAT returned reduced coverage in CpG islands relative to their mean CpG coverage. This pattern was recapitulated around transcript start sites (TSS), where TruSeq was overrepresented, Nanopore and TrueMethyl stayed relatively flat, and EMSeq, MethylSeq, and SPLAT were respectively underrepresented in TSS (Figure 2f).

### Methylation Percentage across Genomic CpGs

After comparing coverage of CpGs, we examined estimates of per-site methylation across assays. As expected, we found methylation percentages to be bimodally distributed with peaks near 0% and 100% methylation. All assays exhibited enrichment for fully methylated regions (Figure 3a), with the exception of Nanopore, which showed underrepresentation of fully methylated regions, a current limitation of its underlying base modification calling method (see methods). For short read approaches, we calculated and corrected for methylation bias (or “mbias”), a measurement of overinflated hypo- or hyper-methylation signal toward the 5’ and 3’ ends of reads. Mbias analysis revealed assay-specific deviation at read ends (Figure 3b). We trimmed bases uniquely for each sample where values began to inflate as recommended by MethylDackel. Mbias analysis also revealed overall methylation trends, with SPLAT and EMSeq tending to have the highest average methylation across reads, while TrueMethyl had the lowest among short read protocols, and TruSeq was the most variably methylated per base across reads.

**Figure 3:**
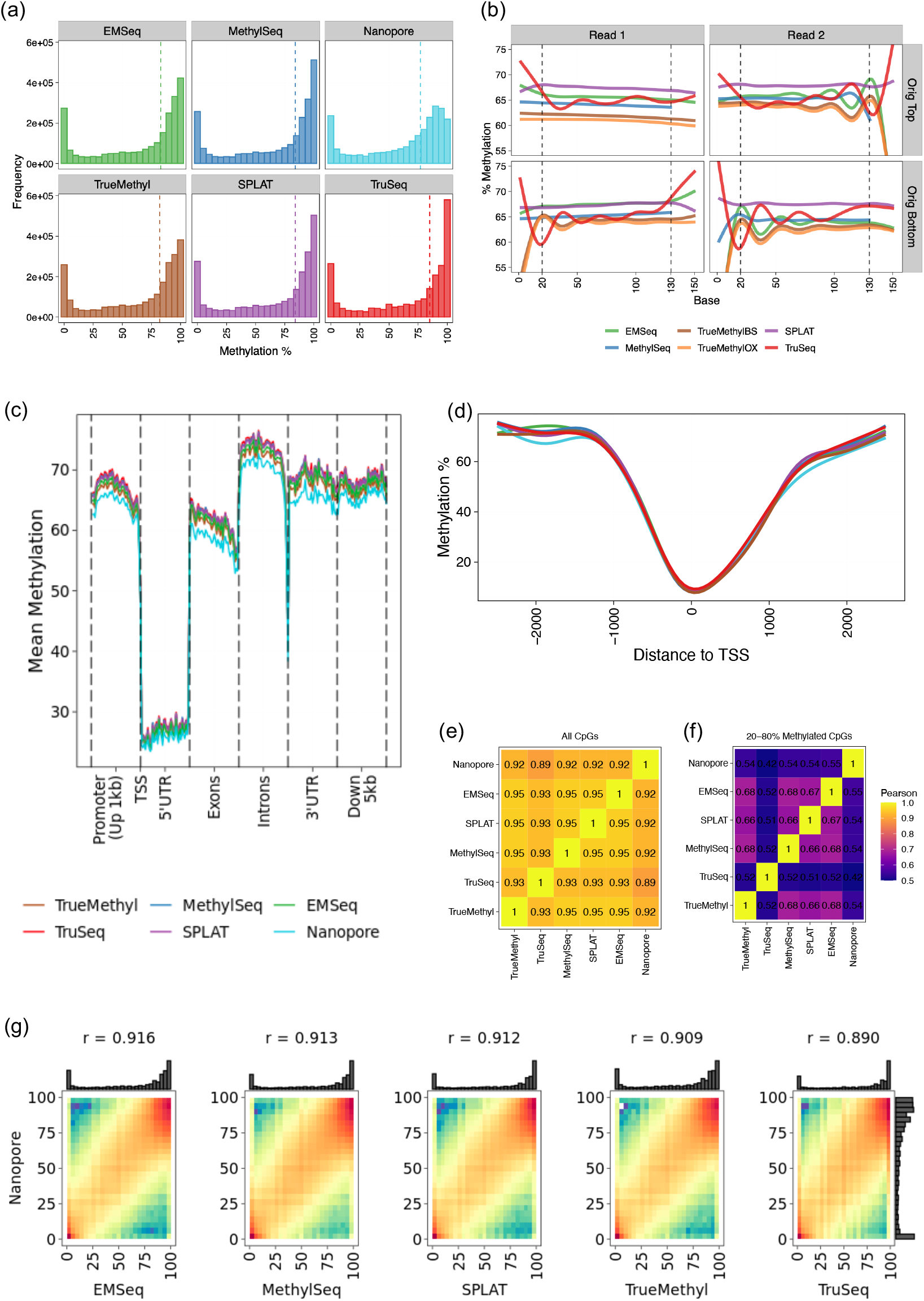
Estimates of methylation per CpG across the genome for HG002. All samples visualized here were downsampled to 20X mean coverage per CpG. (a) Methylation percentage distributons per assay. (b) Methylation bias (mbias) plots showing mean methylation per base for short read assays (Nanopore excluded here). Dotted lines indicate recommended cutoffs for methylation calling for these data. Original Top/bottom refer to mappings to bisulfite-converted strands in the reference genome. (c) Metagene plot showing mean methylation across genomic feature per assay. Promoter regions span 1kb upstream of transcript start sites (TSS). (d) Mean methylation curves surrounding TSS across all genes. (e) Pearson correlation matrix of genome-wide methylation estimates. (f) Pearson correlation matrix of methylation estimates for sites where methylation was estimated to be between 20-80%. (g) Methylation percentage correlation between Oxford Nanopore and all other assays. Pearson correlation values shown on top. Marginal histograms show methylation curves per assay.

We next assigned genomic features to each CpG and summarized methylation across regions in a meta-gene plot (Figure 3c). As expected, we found that methylation levels dropped significantly at TSS and then rose again beyond the 5’UTR in all assays. As detected in the global analysis, methylation captured by Nanopore was lower than by short read assays. Nevertheless, all assays including Nanopore showed highly similar methylation profiles around transcript start sites (TSS) genome-wide (Figure 3d). Correlation of methylation values across genome-wide CpGs was very high (Figure 3e). However, concordance broke down among all assays when restricting to sites with 20-80% methylation, where correlations were as low as r=0.42 between Nanopore and TruSeq (Figure 3f). Therefore the majority of disagreement between assays fell in CpG sites that were either hemimethylated, clonally complex, or undercovered with respect to the global mean. Although short read protocols had higher concordance with one another (r>0.93 for all pair-wise short read comparisons) than with Nanopore estimates, we found that methylation estimation from Nanopore base modification calling was comparable to short read protocols, with Pearson correlation values around r=0.90 for all pairwise comparisons (Figure 3g).

### Family Trio Differential Methylation

Differential methylation was examined at the family trio level. For each methylome assay, we used the replicate-combined methylation calls (including merging bisulfite and oxidative-bisulfite replicates for TrueMethyl) that were normalized to 20X mean coverage.

A total of 2,298,846 CpG sites were present on Chromosome 1 in all six assays (EMSeq, MethylSeq, Nanopore, SPLAT, TrueMethyl, and TruSeq). Coverage levels on HG002 were positively correlated among EMSeq, MethylSeq, and TrueMethyl (Spearman’s *ρ ≥* 0.24). SPLAT coverage was also correlated with these three assays as well as with TruSeq, which was only weakly correlated with any other assay. Nanopore coverage was uncorrelated with that of any other assay. The magnitude of pairwise coverage correlations within each assay varied considerably, with the highest levels observed for TruSeq (0.85 *≤ ρ ≤* 0.86, SPLAT (0.62 *≤ ρ ≤* 0.71), and MethylSeq (0.47 *≤ ρ ≤* 0.48), and the lowest for Nanopore (0.14 *≤ ρ*0.22), EMSeq (0.28 *≤ ρ ≤* 0.31), and TrueMethyl (0.32 *≤ ρ ≤* 0.34).

For each assay, differential methylation analysis was independently conducted at the family level (Ashkenazi Trio HG002-HG004 against the Chinese Trio HG005-HG007). This also included a restriction to sites with 5X coverage in at least two out of three members of each family group, resulting in small data reductions for EMSeq, MethylSeq, Nanopore, SPLAT, and TrueMethyl (3%, 4%, >1%, 4%, and 3%, respectively), and a greater loss for TruSeq (14%). Comparative analysis considered only the 1,928,536 CpG sites that met this criterion for all six assays. To assess consistency in sites identified as differentially methylated (DM) by each assay (DMA), we computed the fraction of DMA sites that were unique to each assay (a pseudo false-positive rate) (Supplementary Table 3). We also computed the total number of DM sites commonly identified by four or more assays (DM4+), which totaled 1.5% of the common sites. We then determined the percentage of DMA sites that were also DM4+ sites (a measure of specificity), as well as the percentage of DM4+ sites that were also DMA sites (a measure of sensitivity).

For EMSeq, 26% of the sites identified as DM were unique to that assay, comparable to MethylSeq (26%) and SPLAT (29%). These three assays were also comparable in the percentage of DM sites that were identified in at least three other assays (36%, 38%, and 35% for EMSeq, MethylSeq, and SPLAT, respectively), and in the percentage of DM sites called by at least three other assays that they also detected (90%, 86%, and 89%, respectively). TrueMethyl detected fewer DM sites overall, with 22% of sites unique to this assay and 42% detected in at least three other assays. However, this did not correspond to a large decline in sensitivity, as 85% of the sites detected by three or more other assays were also identified by TrueMethyl. The smallest number of DM sites was identified in the Nanopore samples, with high specificity (17% unique DMAs and 56% of sites in DM4+) and lower sensitivity, identifying only 51% of the sites identified by four or more other assays. TruSeq, on the other hand, was associated with the largest number of DMA sites and had poor agreement with the other assays, with 43% unique sites, 38% of its sites identified in two or more other platforms, and only 71% of the sites identified by three or more platforms among its DMAs.

Figure 4 illustrates the role of coverage variability for each platform. For each assay, the range between the 5th and 95th percentile of median coverage is shown along the x-axis, while the degree of agreement with other assays for DM sites is shown along the y-axis. We see that agreement declines at higher coverage levels, but this effect is minimal for EMSeq, MethylSeq, Nanopore, and TrueMethyl. Because SPLAT has a more heavy-tailed coverage distribution with stronger sample-to-sample correlations, the impact is more pronounced, while for TruSeq the coverage distribution is extremely diffuse and there is markedly poor agreement with other platforms in its upper coverage percentiles.

**Figure 4:**
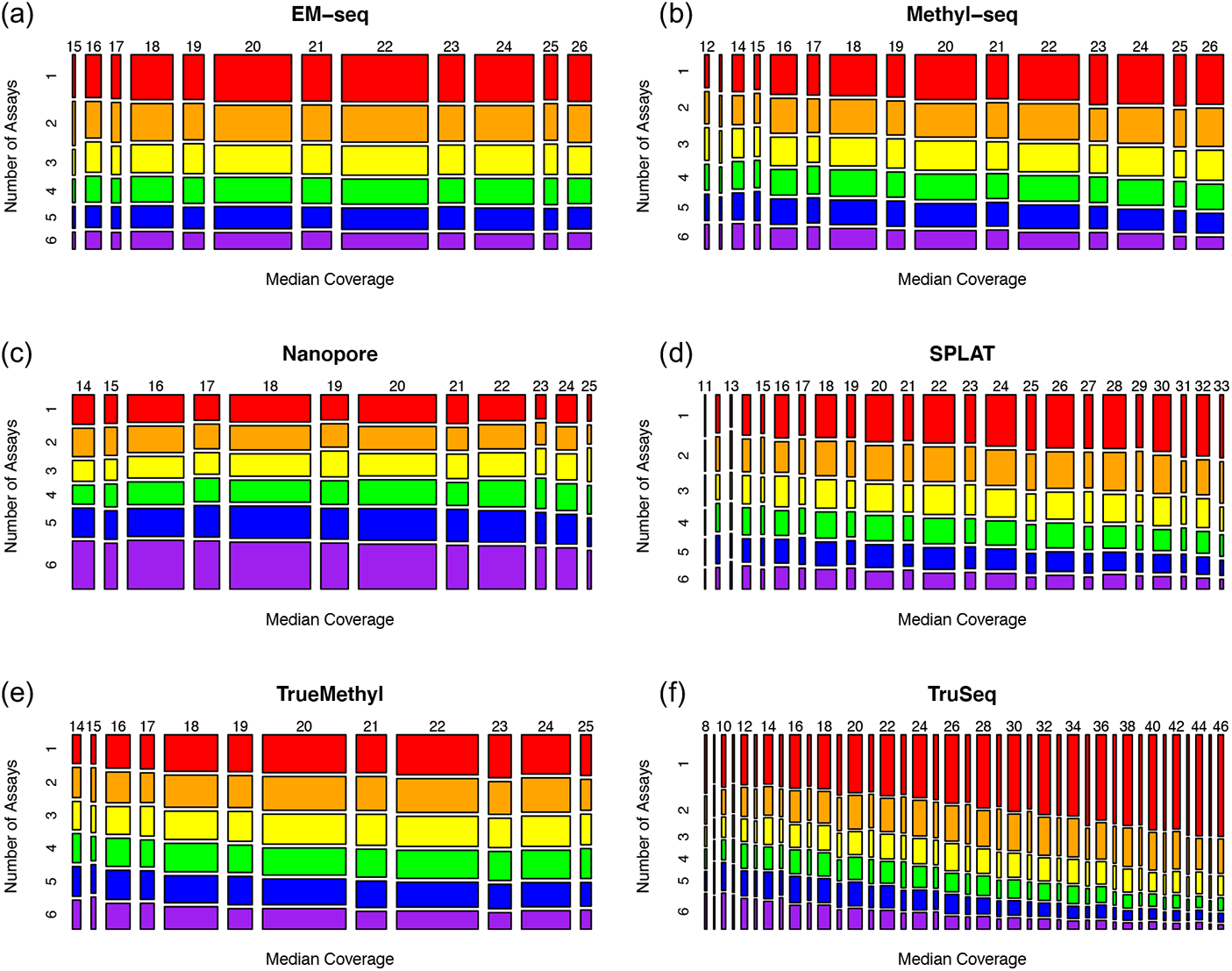
Mosaic plots illustrating agreement between assays for differentially methylated per assay (DMA) sites as coverage levels vary. Rows represent the number of the six assays for which each DMA site was also identified, with values ranging from 1 (indicating no other assays, shaded in red) to 6 (indicating all assays, shaded in purple). Columns indicate the median coverage across HG002-HG007, with values ranging between the 5th and 95th percentiles for each assay.

### Normalization of Array Data

In addition to bisulfite sequencing, microarrays are another commonly used technique to interrogate the epigenome. For each cell line, across three laboratory sites, we generated 3-6 biological or technical replicates with microarray data from the Illumina MethylationEPIC Beadchip (850k array) (Table 1). As a first step before integrating microarray data with the sequencing data, we assessed the performance of different microarray normalization pipelines.

We implemented 26 normalization pipelines with different combinations of between-array and within-array normalization methods. The between-array normalization methods evaluated were no normalization (None), quantile normalization (pQuantile) [18], functional normalization (funnorm) [19], ENmix [20], dasen [21], SeSAMe [22], and Gaussian Mixture Quantile Normalization (GMQN) [23]. The within-array normalization methods evaluated were no normalization (None), Subset-quantile Within Array Normalisation (SWAN) [24], peak-based correction (PBC) [25], and Regression on Correlated Probes (RCP) [26]. All combinations were implemented with the exception of pQuantile + SWAN and SeSAMe + SWAN, which were not possible due to incompatible R object types.

We first performed principal component analysis (PCA) and visually inspected the first two principal components (PCs) for each normalization pipeline (Figure S10). Generally, samples from the same cell line clustered together more tightly after normalization, although a few pipelines (PBC alone, GMQN alone, GMQN + PBC) did not show obvious improvement in replicate clustering. Most pipelines failed to clearly distinguish samples from cell lines HG005 and HG006, the Han Chinese father/son pair, from one another.

A variance partition analysis was used to compute the percentage of methylation variance explained by cell line, lab, or residual variation at each CpG site in each normalized dataset. A superior normalization pipeline would have more variation explained by cell line across the epigenome compared to other pipelines as well as clear clustering of biological and technical replicates.

Funnorm + RCP had the highest median across the epigenome (90.4%), although many pipelines had medians in the 85-90% range (Figure 5a). SeSAMe and RCP performed well (median > 85%) no matter which methods they were combined with. While using RCP or SWAN usually improved performance compared to having no within-array normalization, using PBC for within-array normalization always reduced the median variance explained by cell line. For all downstream analyses, we used the funnorm + RCP normalized microarray data because this pipeline had the highest median variance explained by cell line. Figure 5a shows the full distribution of variance explained by cell line across the epigenome for each normalization pipeline. Most pipelines had a bimodal distribution, so CpG sites typically had almost no variation explained by cell line or nearly 100% of variation explained by cell line.

**Figure 5:**
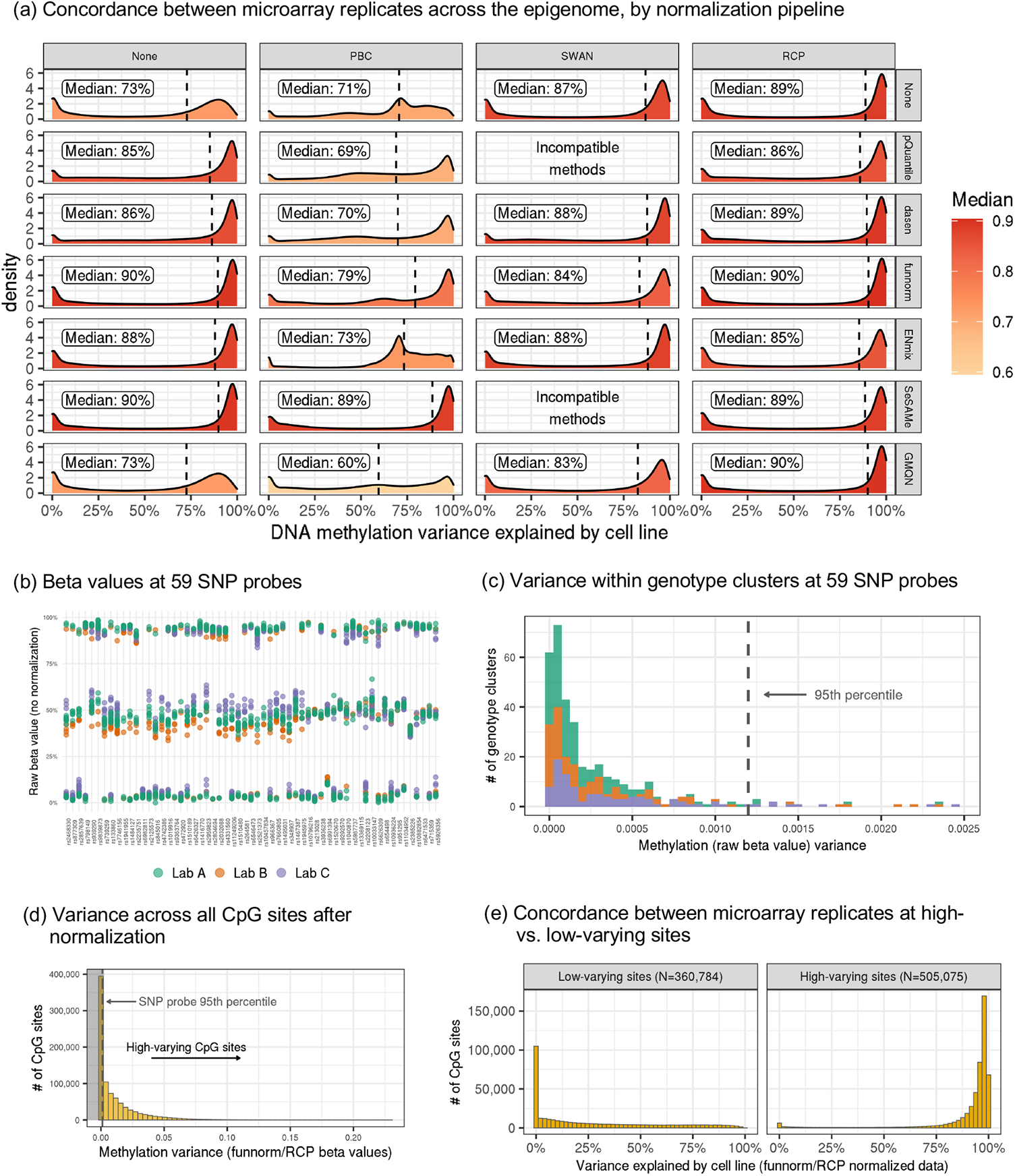
Microarray normalization and low-varying site definition. (a) Densities showing the percentage of DNA methylation variation explained by cell line across the epigenome (N=677,520 overlapping CpG sites) for each normalization method. (b) Raw beta values at each of the 59 SNP probes on the Illumina EPIC arrays, with samples colored by lab. (c) Variance in methylation beta values (no normalization) within each genotype cluster at the 59 SNP probes, separated and colored by lab. The dotted vertical line represents the 95th percentile. (d) Variance in methylation beta values (normalized with funnorm + RCP) across the epigenome. Sites in the shaded area, which have less variation than 95% of SNP probe genotype clusters, are defined as low-varying sites. (e) Percentage of methylation (normalized with funnorm + RCP) variance explained by cell line across the epigenome, stratified by high-varying vs. low-varying sites.

In light of previous work that has shown that microarray data is not reliable for sites with low population variation [27], we investigated whether sites with poor concordance between replicates (% variance explained near 0) overlapped with low-varying sites. We used the 59 SNP probes on the Illumina EPIC array to compute a data-driven threshold for categorizing sites as low varying (Figure 5b-d; see methods for details). We found that nearly all CpG sites in the normalized (funnorm + RCP) microarray data with poor concordance between replicates met our definition of low-varying sites (Figure 5e). This suggests that our data-driven definition of low-varying CpG sites, which can be applied to any Illumina 450k or 850k array dataset, may be useful for filtering out less reliable CpG sites before analysis.

### Normalized Microarray Concordance with Sequencing Data

We performed 6 additional variance paritition analyses, adding samples from one sequencing assay (EM-Seq, MethylSeq, SPLAT, TrueMethyl, TruSeq, or Nanopore) at a time, to evaluate the concordance between microarray and downsampled 20X sequencing data. For each site and each sequencing assay, we estimate the percentage of methylation variance explained by cell line, assay (sequencing or microarray), and residual variation. A higher percentage of variance explained by cell line indicates better agreement with the microarray data. Ternary density plots of the variance explained by cell line, assay, or residual variation show lower concordance between the Nanopore sequencing data and the microarray data than other sequencing assays (Figure 6a). The five other sequencing assays (EMSeq, MethylSeq, SPLAT, TrueMethyl, and TruSeq) have a high density of sites where nearly 100% of the methylation variance in the merged seqeuencing/microarray dataset is explained by cell line. However, for all assays, there is a smaller peak of CpG sites where nearly 100% of the methylation variance is explained by assay, indicating that there were some technical artifacts introduced by assay, but these technical artifacts were not widespread across the epigenome.

**Figure 6:**
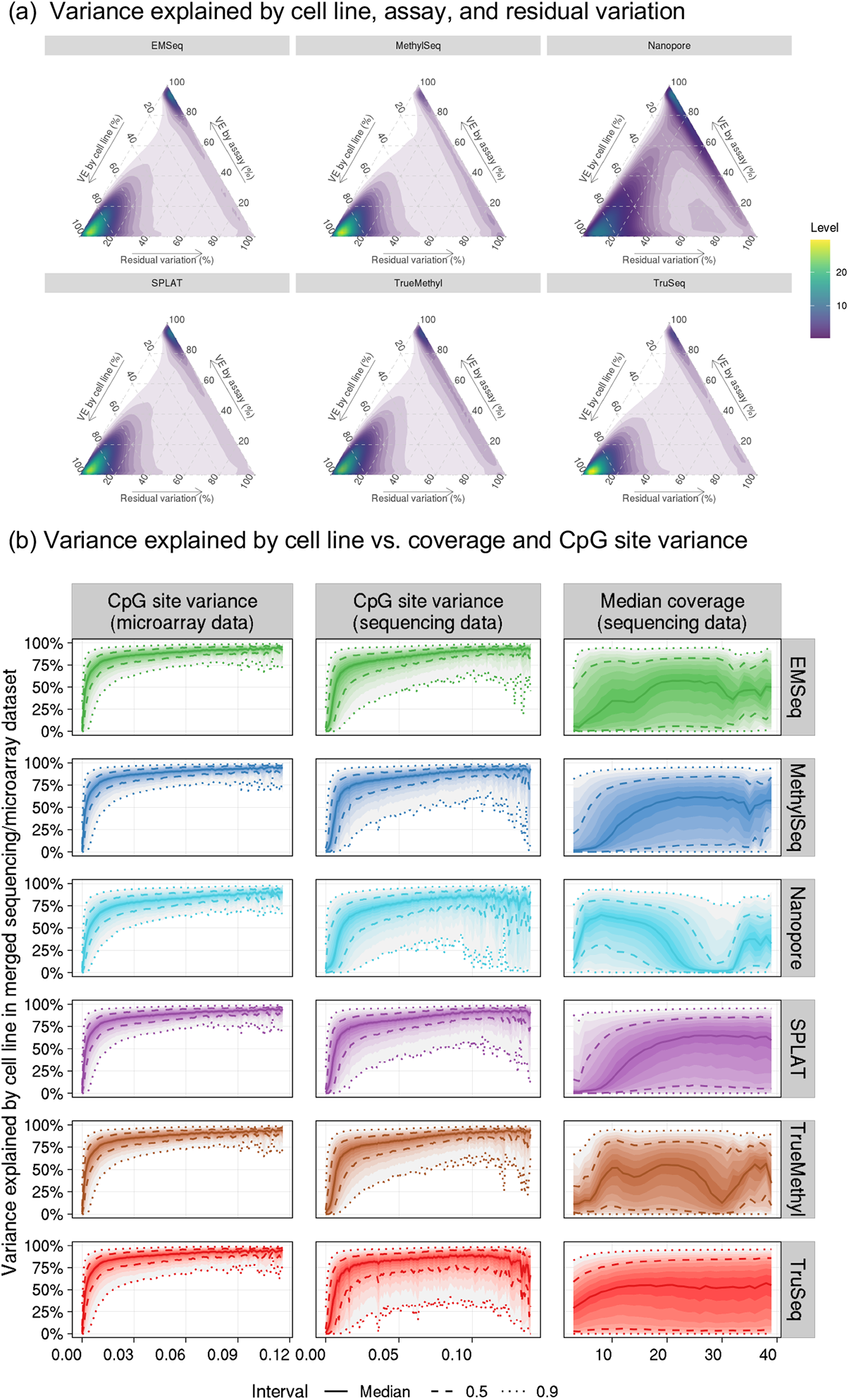
(a) Density plots of sequencing/microarray concordance indicating the percent of variance explained (VE) by cell line, assay (sequencing or microarray), and residual variation for 841,833 CpG sites with complete information in all assays. (b) Distribution of percent variance explained by cell line in the sequencing/microarray variance partition analysis as a function of beta value variance (binwidth=0.001) and median coverage (binwidth=1) at each CpG site. 90% of the y-axis values fall between the outermost dotted lines for each bin along the x-axis.

We investigated what was driving poor concordance between assays at this subset of CpG sites and found a strong, non-linear relationship between the amount of variability at a CpG site and concordance (Figure 6b). The non-linear relationship between CpG site variance in the microarray data and concordance between assays indicates that there is a minimum amount of population variance needed for reproducibility, but beyond this threshold more variation does not improve concordance. This confirms our proposed approach of estimating technical noise from the SNPs on the array to create a binary “low-varying” or “high-varying” classification for CpG sites.

Because each cell line had 3-6 microarray replicates and only one (merged replicate) sequencing sample, these results are largely driven by the microarray data and the estimates of the percentage of variation explained by cell line (vs. assay) are likely biased upward by this. Visual inspection of the joint distribution of microarray and sequencing beta values for all HG002 replicates (with sequencing replicates from the same lab merged) shows that there is substantial technical noise in the data when comparing any two assays (Figure S11). For the same assay in two different labs, we see much better concordance between HG002 beta values with microarrays than with EMSeq.

### Differential Methylation in Microarray Sites

We took differentially methylated regions between family groups (see above) and restricted them to sites captured by the Illumina MethylationEPIC Beadchip (850k array) (see above). Of the 82,013 probes on the array that map to regions on Chromosome 1, 81,456 sites (99.3%) were detected at high depth by all six sequencing assays. Of these, the number of differentially methylated assays (DMAs) ranged from 1,027 (Nanopore) to 4,267 (TruSeq). For EMSeq, MethylSeq, Nanopore, and TrueMethyl, over 99% of these DMA had estimated percent methylation difference (PMD) of 20% or greater between the family groups, while 95% and 80% of DMAs met this criterion for SPLAT and TruSeq, respectively.

To analyze concordance between the sequencing-based and array results, we computed the proportion of these DMAs for which a corresponding difference of at least 20% was observed for the arrays, with these array PMDs estimated via ANOVA models with random intercepts for each genome. As illustrated Supplementary Table 4, the overall agreement was comparable for four of the six methods with values ranging from 55.5% (EMSeq) to 60.0% (TrueMethyl), with a higher level of 67.0% for Nanopore and a lower level of 49.6% for TruSeq. However, among the 4,137 sites with array |PMD|>0.2, only 16.6% were Nanopore DMAs in comparison to 42-44% for all other assays, suggesting high precision but lower sensitivity for this assay.

## Discussion

The EpiQC study provides a comprehensive epigenetic benchmarking resource using human cell lines established by the Genome in a Bottle Consortium as reference materials to advance genomics research. We provide datasets for a broad range of methylome sequencing assays, including short-read whole genome bisulfite sequencing (WGBS) and enzymatic deamination (EMSeq), and native 5-methylcytosine calling using Oxford Nanopore long read sequencing. We also provided data from targeted approaches, including reduced representation bisulfite sequencing (Methyl Capture EPIC) and the Illumina Infinium MethylationEPIC 850k array. While most of the published and/or commercialized assays have been tested with some standard sample (e.g. GM12878), the sample used to benchmark each assay was drawn from different DNA aliquots, extracted from cells grown at different passage, and potentially grown in different media. Here, aliquots of the same gDNA were distributed across multiple laboratories, and used for all data generated. To remove additional variability, all libraries were sequenced on multiple flow cells of one Illumina NovaSeq 6000 (then a third flow cell on the same instrument type). For all assays, libraries were produced in duplicates, providing both inter- and intra-assay datasets.

Benchmarking whole methylome sequencing technologies is important for determining which method will achieve the best performance, and to provide recommendations and standards for experimental design within future studies. Large projects such as the NIH Roadmap Epigenomics Project [28] the International Human Epigenome Consortium [29], and the Cancer Genome Atlas [30] have produced, compiled, and analyzed a vast amount of WGBS data comprising tissues and cell lines from normal and neoplastic tissues. Building upon these previous works, our study encompasses an up-to-date range of commonly used whole methylome assays as well as emerging methods such as enzymatic methylation and native 5mC calling from long read technologies, and provides data across 7 different reference material cell lines, providing a comprehensive examination of DNA methylation analysis methods.

We found that the library prepration method of choice and parameters used within each protocol had an outsized impact on data quality and biological inference. Libraries with longer inserts benefitted from less adapter contamination, fewer dovetailing (overlapping) reads, and fewer low quality bases, which increased mapping efficiency and mean coverage per CpG. This is particularly impactful when one chooses to employ a cost-effective sequencing on an Illumina system with paired-end 150 bp reads, as was done within this study. This sequencing scheme resulted in a highly variable depth of coverage per library preparation. While imbalanced pools may account for some of the difference, library preparation methods had the biggest impact. Except for TruSeq, all the other library preparations start with shearing of the gDNA. For the other bisulfite-dependent protocols, the DNA fragments range between 200-400, whereas EMSeq allows for longer fragments (550bp). TruSeq libraries tend to have short (130 bp) insert sizes and are therefore more suitable for 75 bp paired-end read lengths. To overcome the impact of imbalanced sequence depth, this study provides robust recommendations for downsampling across sequencing types, showing both how different downsampling schemes (i.e. at the BAM level or at the methylation bedGraph level) are comparable, and how downsampled datasets can be directly compared to one another to assess the performance of the assays themselves.

The methods that have proven to have greater genome-wide evenness of coverage, namely Accel-NGS MethylSeq [15], SPLAT [6], and TrueMethyl [31] tend to have longer insert sizes (200–300 bp), fewer PCR duplicates (down to a few percent, depending on sequencing platform), and high mapping efficiencies (>75%). The SPLAT libraries herein had shorter insert sizes than desired due to the use of 400 bp Covaris shearing prior to library preparation. To achieve insert sizes of >=300bp, the SPLAT authors now recommend using DNA fragmented to 500-600 bp as input and to perform final library purification at 0.8x AMPure ratio to remove shorter fragments. The same recommendation may also improve the insert size for MethylSeq and TrueMethyl protocols. SPLAT is the only method in our evaluation that is not commercial/kit-based and could be comparatively ~10x cheaper per library [6]. This can be important when considering the sample preparation cost alongside sequencing costs.

NEB’s EM-Seq protocol [32] compares favorably to the bisulfite sequencing-based approaches analyzed herein. In almost all comparisons EM-Seq libraries captures more CpG sites at equal or better coverage. We also show that the methylation signal achieved by native base modification detection from Oxford Nanopore long read sequencing is highly comparable to short read bisulfite- and enzymatic-methylation sequencing, with average Pearson correlation values of r=0.90 for CpG methylation concordance. Moreover, Nanopore can detect a significant number of sites that short read assays miss, many of which occur in promoter and exonic regions that are potentially of biological significance.

Beyond library preparation, the use of algorithmic tools has an impact on the performance of each methylome assay. Asymmetrical C-T distributions between DNA strands and reduced sequence complexity make epigenetic sequence alignment different from regular DNA processing. We compared common methylation processing piplines and compared their mapping efficiencies, depth of coverage achieved per CpG, and computational time to run, and observed bwa-meth to provide the best performance when considering all of these factors. Notably, BitMapperBS was significantly faster and not far behind bwa-meth in terms of mapping efficiency and CpG coverage.

Another important parameter is the amount of data retained from a WGBS experiment following adapter and quality trimming, mapping and deduplication. Here, we show the effects of each mapping step on each methylome assay (Figure S4), and how reads are filtered along each step, including the estimated number of reads required to achieve a certain mean coverage per CpG (Table 2). Similarly, previous studies [5, 15] have implemented a metric to estimate the efficiency of WGBS genome coverage by determining the raw library size (number of PE 150 bp reads prior to filtering) required to achieve at least 30x coverage of 50% or more of the genome. We propose a modified version of the calculation proposed by Zhou and collegues, deriving the number of PE150 bp reads needed to achieve 20x average CpG coverage for a library, as this metric directly relates back to the CpG sites whose methylation levels will be interrogated. We also calculate useable bases, reflecting the total bases used for methylation estimation out of the total bases sequenced for per library. Adoption of such metrics will make it significantly easier to compare and contrast results from different methods.

Choice of computational algorithms is equally important in analyzing methylation microarray data. In this study, we compared 26 different normalization pipelines. Many algorithms (SWAN, RCP, pQuantile, dasen, funnorm, ENmix, SeSAMe) generally performed well in this dataset, clustering replicates from the same cell line together while preserving differences between cell lines. Given the comparable performance of these methods, the best normalization pipeline will depend on the needs of individual studies. For instance, cohorts with multiple tissues may want to use the multi-tissue extension of funnorm, funTooNorm[33], and cohorts with very large sample sizes may want to use SeSAMe[22], which is the only single-sample normalization method we evaluated. All pipelines performed poorly at sites with low population variance, confirming previous work [27]. We propose using the SNPs on the 850k array to calculate a data-driven threshold for classifying and filtering out low-varying sites before analysis. Previously published associations at sites with low population variation, which can also often be identified by their extreme (<5% or >95%) median methylation values[27], should be interpreted with caution. Aditionally, our data from EMSeq and microarray replicates across different labs Figure S11 support previous findings that the Illumina 850k array was more reproducible than TruSeq across paired technical replicates from 4 cord blood samples [34]. We conclude that overall, microarrays are a good option for researchers who are comfortable with a targeted assay.

One final caveat for the data within this study is our use of high quality DNA from EBV-immortalized, B-lymphoblastoid cell lines. Using this highly controlled input, the methods examined within this study produced mostly comparable data. However, the performance of each kit may be more variable on less optimal input DNA (lower input, more highly fragmented, etc.) that mirrors real clinical samples more closely. The optimal data herein should serve as a launch point for future studies of more realistic inputs.

## Methods

### Genomic DNA

The samples in this study comprise genomic DNA (gDNA) from seven EBV-immortalized B-lymphoblastoid cell lines designated as reference samples by the National Institute of Standards and Technolog (NIST) Genome in a Bottle Consortium (see https://www.coriell.org/1/NIGMS/Collections/NIST-Reference-Materials). The NA12878 (HG001) cell line was selected as it is the most commonly used reference for benchmarking or generation of genomics datasets. Additionally, six cell lines representing two trios from the Personal Genome Project, which are consented for commercial redistribution, were also included. The HG002/3/4 samples were provided by a son/father/mother trio of Ashkenazi Jewish ancestry, and the HG005/6/7 come from a Han Chinese son/father/mother trio.

For each reference cell line, 100 ug genomic DNA (gDNA) was purchased from the Coriell Institute for Medical Research, along with viable cell lines for later growth and distribution. The gDNA was quantitated using Qubit Broad Range dsDNA kit and an aliquot from reference sample gDNA was distributed to six independent laboratories for NGS library preparation or microarray analysis.

### NGS Library Preparation

#### Enzymatic Methyl-Seq (EMSeq)

EMSeq libraries were prepared at two different laboratories using slightly altering protocols. At Lab1, genomic DNA was spiked in with 2 ng unmethylated lambda as well as 0.1 ng CpG methylated pUC19, and was then fragmented to 500 bp using a Covaris S2 (200 cycles per burst, 10% duty-cycle, intensity of 5 and treatment time of 50 seconds). At Lab2, genomic DNA was fragmented to 450 bp using Covaris 130uL. While all replicates of HG001-004 were created using 100ng of DNA, both labs created replicates of HG005-007 using 100ng, 50ng, and 10ng of DNA in order to test the effects of input concentration. EM-seq libraries from both laboratories were prepared using the NEBNext Enzymatic Methyl-seq (E7120, NEB) kit following manufacturer’s instructions. Final libraries were amplified with NEBNext Q5U polymerase using 4 PCR cycles for 100 ng, 5 cycles for 50 ng and 7 cycles for 10 ng inputs. Libraries were quality controlled on a TapeStation 2200 HSD1000.

#### Swift Biosciences Accel-NGS Methyl-Seq (MethylSeq)

Libraries were prepared according to manufacturer’s instructions (Swift) using dual-indexing primers. Briefly, 100ng of genomic DNA was spiked in with 1% unmethylated Lambda gDNA, and fragmented to 350 bp (Covaris S220, 200 cycles per burst, 5% duty-factor, 175W peak displayed power, duration of 50 seconds). Bisulfite conversion was performed using EZ DNA Methylation-Gold kit (Zymo Research). Adaptase was used to ligate adapters to the 3’ end of the bisulfite-converted DNA, followed by primer extension, second strand synthesis, and ligation of adapter sequences at its 3’ end. The libraries were amplified for a total of 6 rounds using the Enzyme R3 provided with the kit. Libraries were quality controlled on a TapeStation 2200 HSD1000.

#### SPlinted Ligation Adapter Tagging (SPLAT)

100ng gDNA was fragmented to 400 bp (Covaris E220, 200 cycles per burst, 10% duty factor, 140 peak incident power PIP, 55s treatment time). Bisulfite conversion was performed using the EZ DNA Methylation-Gold kit (Zymo Research). SPLAT libraries were constructed as described previously [6]. Briefly, adapters with a protruding random hexamer were ligated at the 3’ end and 5’ end of single-stranded DNA in consecutive reactions. The resulting libraries were amplified with 4 PCR cycles using KAPA HiFi Uracil+ PCR enzyme (Roche). Libraries were quality controlled on a TapeStation 2200 HSD1000.

#### NuGEN TrueMethyl oxBS-Seq (TrueMethyl)

200 ng of genomic DNA was spiked with 1% unmethylated Lambda gDNA and fragmented to 400 bp (Covaris S220, 10% duty-factor, 140W peak incident power, 200 cycles per burst, duration of 55 seconds). Fragmented DNA was processed for end-repair, A-tailing, and ligation using NEB’s methylated hairpin adapter. Ligation was performed at 16°C overnight in a thermocycler. The USER enzyme reaction was performed the next morning, according to the manufacturer’s protocol, and the adapter-ligated DNA cleaned up using 1.2:1 Ampure XP bead:ligated DNA ratio. Each ligation was then split into 2 aliquots to perform oxidation + bisulfite conversion or mock (water) + bisulfite conversion according to the OxBS module instructions (Tecan/NuGen). PCR amplification was performed using NEB’s dual-indexing primers and KAPA Uracil+ HiFi enzyme for a total of 10 cycles. Libraries were quality controlled on a TapeStation 2200 HSD1000.

#### Illumina TruSeq DNA Methylation (TruSeq)

100ng of genomic DNA was bisulfite converted using EZ DNA Methylation-Gold Kit (Zymo Research). Sequencing libraries were prepared according to the manufacturer’s protocol (Illumina). Briefly, the bisulfite-converted DNA was first primed by random hexamers containing a tag sequence on its 5’ end. Next, the bottom strand was extended and a 3’ end oligo added. The libraries were amplified with 10 PCR cycles using the FailSafe PCR enzyme (Illumina/Epicentre). Libraries were quality controlled on a TapeStation 2200 HSD1000.

#### Illumina Methyl Capture EPIC

500ng of genomic DNA was prepared according to the manufacturer’s protocol (Illumina), including a spike-in of 2 ng of unmethylated lambda. Briefly, the genomic DNA was fragmented to 200 bp using a Covaris S220 (10% duty-cycle, 175W peak incident power, 200 cycles per burst, duration of 360 seconds). The fragmented DNA was next purified using AMpure XP beads, end repaired and A-tailed, before ligation of single index adapters with methylated cytosines. Libraries cleaned using AMpure XP beads, then pooled in 3- and 4-plex. The pools were denatured to single stranded DNA before hybridization to the RNA baits provided with the kit. After cleanups of the hybridizations according to the manufacturer’s protocol, the captured strands were process for library amplification by PCR using KAPA Uracil+ HiFi enzyme (Roche) and TrueSeq primers included in the kit. Libraries were quality controlled on a TapeStation 2200 HSD1000..

#### Oxford Nanopore Library Preparation

Genomic DNA was quantified using a Qubit 4 Fluorometer (ThermoFisher Q33238) and libraries were prepaird using a Ligation Sequencing Kit (SQK-LSK109, Oxford Nanopre Technologies). Briefly, 1000ng of genomic DNA was end-repaired and dA-tailed using the NEBNext End Repair/dA-tailing module, and then sequencing adapters were ligated. DNA fragments below 4kb were removed using the long fragment wash protocol option according to the manufacturer’s protocol.

### EPIC Microarrays

#### Illumina Infinium MethylationEPIC BeadChip (850k array)

Bisulfite conversion was performed using the EZ DNA Methylation Kit (Zymo Research) with 250 ng of DNA per sample. The bisulfite converted DNA was eluted in 15 µl according to the manufacturer’s protocol, evaporated to a volume of <4 µl, and used for methylation analysis on the 850k array according to the manufacturer’s protocol (Illumina).

Microarray experiments were run at three different labs, denoted Lab A, B, and C to distinguish them from the sequencing labs (Lab 1 and Lab 2). The resulting dataset contains 30 samples, with each of the seven cell lines (HG001-HG007) having between three and six replicates (biological or technical). Two technical replicates were generated for each cell line at lab A, one replicate from each cell line was generated at lab B, and three technical replicates were generated for the Han Chinese family trio cell lines (HG005-HG007) at lab C.

### LC-MSMS Quantification

#### LC-MS/MS quantification of SmC and ShmC

Genomic DNA from HG001-007 cell lines was used for the analysis. Samples were digested into nucleosides using Nucleoside digestion mix (M0649S, New England Biolabs) following manufacturers protocol. Briefly, 200 ng of each sample was digested in a total volume of 20 µl using 1 µl of the digestion mix. Samples were incubated at 37°C for 2 hours.

LC-MS/MS analysis was performed using two biological duplicates and two technical duplicates by injecting digested DNA on an Agilent 1290 UHPLC equipped with a G4212A diode array detector and a 6490A Triple Quadrupole Mass Detector operating in the positive electrospray ionization mode (+ESI). UHPLC was performed on a Waters XSelect HSS T3 XP column (2.1 × 100 mm, 2.5 µm) using a gradient mobile phase consisting of 10 mM aqueous ammonium formate (pH 4.4) and methanol. Dynamic multiple reaction monitoring (DMRM) mode was employed for the acquisition of MS data. Each nucleoside was identified in the extracted chromatogram associated with its specific MS/MS transition: dC [M+H]+ at m/z 228-112, 5mC [M+H]+ at m/z 242-126, and 5hmC [M+H]+ at m/z 258-142. External calibration curves with known amounts of the nucleosides were used to calculate their ratios within the analyzed samples.

### DNA Sequencing

#### Illumina sequencing

The short-read sequencing libraries were collected from participating laboratories and sequenced centrally at two sequencing centers. Libraries were pooled by library type in high concentration equimolar stock pools (4 nM). After pooling, bead-based clean-up was performed to remove peaks <200 bp. The cleaned stock pools were quantified on an Agilent Bioanalyzer using High sensitivity DNA chip and subsequently diluted to 1.5 nM prior to sequencing on Illumina NovaSeq 6000 S4 flowcells PE150 read-length to a targeted minimum per replicate CG coverage of 20x. Base calling was performed using RTA v3.4.4. Additional details about the sequencing parameters can be found in the Supplementary Materials and Methods.

#### Oxford Nanopore Sequencing

The Nanopore libraries were run simultaneously on seven FLO-PRO002 flowcells for 64 hours on a PromethION Beta device to maximize yield. FAST5 files were generated using default parameters within MinKNOW on the PromethION machine. Base calls and base modification calls were generated using Megalodon v2.2.9 (https://nanoporetech.github.io/megalodon/) with guppy v4.2.2 (https://community.nanoporetech.com/downloads/guppy) as the basecaller backend. The MinION DNA R9.4.1 5mC configuration file from the Rerio database (https://github.com/nanoporetech/rerio) was used as the base modification model. The MinION model was chosen because it maintained more consistent peaks at 0% and 100% methylation as compared to the PromethION model.

### Data Quality Control

FastQC (https://www.bioinformatics.babraham.ac.uk/projects/fastqc/) was used to evaluate the quality of sequencing data, including base qualities, GC content, adapter content, and overrepresentation analysis. Adapter sequences were trimmed using FASTP [35] with a minimum length of two bases, quality filtering disabled, and forced poly-G trimming. The data generated using the Swift Methyl-Seq kit were further trimmed for an additional 10bp on the 3’ end of R1 and 10bp on the 5’ end of R2 to remove Adaptase sequence introduced during library prepartion.

### Alignment and Methylation Calling

Alignment comparison was conducted on sample HG002. All short read WGBS libraries were aligned to the human reference genome (build GRCh38) with additional contigs included representing bisulfite controls spiked within pooled libraries, including lambda, T4, and Xp12 phages, as well as cloning vector plasmid pUC19. The Epstein-Barr Virus (EBV) sequence was also included as a decoy contig to account for use of EBV to immortalize B-lymphocytic cell lines.

#### BISMARK

Adapter-trimmed reads were aligned using two parallel instances of BISMARK v0.23.0 (https://github.com/FelixKr per replicate and bowtie2 (http://bowtie-bio.sourceforge.net/bowtie2/index.shtml) as the read aligner. BAM files were position sorted using sambamba sort (https://lomereiter.github.io/sambamba/) and deduplicated using deduplicate_bismark with default paramters. Methylation was called using bismark_methylation_extractor using 2 multicore instances and default parameters and strands were merged into dinucleotdie contexts using MethylDackel (https://github.com/dpryan79/MethylDackel) mergeContext.

#### BitMapperBS

Alignment was run using default parameters within BitMapperBS v1.0.2.2 on adapter-trimmed FASTQs and the resulting BAMs were position sorted using sambamba sort. Alignments were deduplicated using Picard MarkDuplicates (https://broadinstitute.github.io/picard). Methylation was extracted using MethylDackel extract and strands were merged into dinucleotide context using MethylDackel mergeContext.

#### BSSeeker2

Adapter-trimmed reads were aligned across four threads within BSSeeker2 using bowtie2 as the aligner per user guide recommendation. Alignments were sorted using sambamba sort and deduplicated using Picard MarkDuplicates. Methylation was called within bs_seeker2-call_methylation and strands were merged into dinucleotdie contexts using MethylDackel mergeContext.

#### bwa-meth

Adapter-trimmed reads were aligned using bwa-meth v0.2.1 with default parameters and converted into BAM format using sambamba view. Alignmens were then position sorted with sambamba sort and deduplicated using Picard MarkDuplicates. Methylation was called with MethylDackel extract and strands were merged into dinucleotdie contexts using MethylDackel mergeContext

#### gemBS

gemBS v3.2.0 (https://github.com/heathsc/gemBS) requires two set-up files to enable analysis. The first file is a metadata sheet, in which sample barcodes were provided in assay/lab/genome/replicate format (e.g. EMSeq_LAB01_HG001_REP01). The second file is a configuration sheet, in which default parameters were applied, including MAPQ threshold of 10, base quality threshold of 13, reference bias of 2, 5’ trim of 5bp, 3’ trim of 0bp, removing improper pairs, marking duplicate reads, diploid alignment, auto conversion, and all files generated (CpG, non-CpG, bedMethyl, and bigWig). These files were fed into gemBS which uses GEM3 for alignment and BScall for methylation calling.

### Downsampling Methylation Calls

The 5-methylcytosine bedGraph files generated by the bwa-meth aligner (see results for rationale to proceed with bwa-meth calls for secondary analyses) were normalized such that each call set had a given mean global coverage per CpG. In order to maximize coverage per library, all technical replicates were combined per library type per cell line per laboratory (e.g., all replicates for EM-Seq HG002 from Laboratory 1 were combined) by summing up the methylated and unmethylated counts per CpG site. Next, counts along the positive and negative strands were merged in order to produce one value per CpG dinucleotide using MethylDackel mergeContext. The resulting replicate-CpG-merged bedgraphs were downsampled using https://github.com/nebiolabs/methylation_tools/ downsample_methylKit.py where a fraction of counts kept corresponding to the desired downsampling depth.

To compare downsampling from mapped reads (BAM files) in comparison to bedGraph files, the BAM files from all replicates representing EMSeq HG006 (Lab 1) were respectively merged using samtools merge. The merged BAMs were then downsampled using samtools view using the *−s* parameter, calculating the fraction of reads necessary to achieve the desired mean coverage per BAM. Methylation was called on these BAM files using the same methodology as above. The strands were merged by CpG dinucleotide using MethylDackel merge context, creating one methylation call per CpG site. The procedure is outlined in Figure S5.

### Differential Methylation

Differential methylation between the two family groups (Ashkenazi Jewish Trio, HG002-HG004 vs Chinese Han Trio, HG005-HG007) was assessed at each site on Chromosome 1 for which at least two samples per group were covered by 5 or more reads. Following aggregation of replicates, strand merging, and down-sampling to mean 20X coverage, analysis was independently conducted via logistic region for each of six platforms (Methyl-seq, EM-seq, Nanopore, TruSeq, SPLAT, and TrueMethyl) using the standard “glm” function in R. *p*-values were adjusted using the Benjamini-Hochberg correction and adjusted values < 0.05 were considered statistically significant. Comparisons among platforms considered only sites that were present for all assays.

### Microarray Normalization

Microarray normalization methods were divided into two broad categories: between-array normalization and within-array normalization. Between-array normalization is used to reduce technical variation while preserving biological variation between samples, while within-array normalization is used to correct for the two different probe designs on the Illumina methylation arrays, which have been observed to have different dynamic ranges [25]. The between-array normalization methods evaluated were pQuantile [18], funnorm [19], ENmix [20], dasen [21], SeSAMe [22], and GMQN [23]. We implemented all possible combinations of between-array and within-array normalization methods as well as each method individually. Samples from all 3 labs were normalized together as one joint dataset.

In order to evaluate the performance of each pipeline, all 30 microarray samples from 3 labs were pooled together in a variance partition analysis [36]. For each pipeline and at each CpG site, the percentage of variation in DNA methylation beta values explained by cell line and lab was calculated. Additionally, we performed principal components analysis (PCA) and visually inspeced clustering of technical and biological replicates across all normalization pipelines.

After normalization, we used the 59 SNP probes on the 850k array, meant to identify sample swaps [37], to define a data-driven classification of low-varying sites. Previous studies have found that low-varying sites have poor reproducibility on the Illumina arrays [27] and have suggested data-driven probe filtering using technical replicates [38, 39] or beta value ranges [27]. However, not all studies have technical replicates, and previously proposed beta value range cutoffs for one experiment may not be generalizable to another experiment. We first called genotype clusters based on the beta values at each of the 59 SNP probe within each of the 3 different labs (**??**b). Although we used a naïve approach for calling genotypes (<25% methylation=cluster 1, 25-50% methylation = cluster 2, >75% methylation = cluster 3), which was sufficient for the clear separation in our dataset (**??**b), more sophisticated methods [40] can be used for datasets with less clear separation and/or outlier values. In theory, because these 59 SNP probes are meant to measure genotypes, cell lines with the same genotype should have exactly the same readout in an experiment without any technical noise. Therefore, we can use variance within genotype clusters from the same experiment as a measure of technical noise and determine the minimum population variation needed to exceed the observed technical variation. Within each of the 3 labs, we calculated methylation variance at each SNP probe within each genotype cluster, giving us a distribution of observed technical noise (**??**c). To avoid being overly conservative due to outlier values at these 59 SNP probes, we use the 95th percentile of these genotype cluster variances as the threshold for defining low-varying sites (**??**c-d).

### Sequencing Performance in Micorarray Sites

Variance partition analyses [36] were used to compare the microarray and downsampled sequencing datasets and assess concordance between microarray and sequencing assays. Each of the variance partition analyses included all microarray replicates, normalized with funnorm + RCP, and one sequencing sample per cell line with all replicates merged. The percent of variation in DNA methylation explained by cell line, assay (sequencing or microarray), and residual variation was calculated at each CpG site. This produced 6 sets of results, one per sequencing assay. The percentage of variation explained by cell line at each site was used as a measure of cross-platform concordance between each sequencing platform and the microarray data. The variance partition results presented are restricted to CpG sites that were measured in all 7 cell lines across all 7 assays (N=841,883) to ensure a fair comparison.

## Data Availability

All data sequenced for this study is available within SRA under accession number SRR8324451. All code used to process data and generate files is publicly available on Github at https://github.com/Molmed/epiqc.

## Acknowledgments

J.N, A.L, U.L, T.A and A.R are supported by grants from the Swedish Research Council (2017-00630 / 2019-01976). I.I.C, R.R, and C.R.A are supported by ISCIII, project number PI18/00050. T.G and Y.P.D are supported by NIH Grants 5P30GM114737, P20GM103466, U54 MD007584, and 2U54MD007601. The genomic work carried out at the Loma Linda University Center for Genomics was funded in part by the National Institutes of Health (NIH) grant S10OD019960 (CW). This project is partially supported by AHA grant 18IPA34170301 (CW).

## Disclaimer

The views presented in this article do not necessarily reflect those of the U.S. Food and Drug Administration. Any mention of commercial products is for clarification and is not intended as an endorsement.

## Author Contributions

C.E.M, Y.W, Y.D, J.M.G, C.W, M.S, M.N, C.S, A.M, J.W.D, W.X, H.H, B.N, and W.T conceived of and designed the study. A.R, U.L, D.B, A.A, G.G, J.I, F.W, V.K.C.P, L.W, C.L, Z.C, Z.Y, J.L, X.Y, H.W, S.G, and D.B.M prepared sequencing libraries. V.K.C.P and L.W pooled and sequenced the libraries. T.A, R.R, C.R.A, I.I.C, T.G, Y.P.D, and M.N generated microarrays. J.F, A.L, J.N, B.W.L, M.L, M.A.C, C.R.A, T.G, C.L, K.P, R.C, S.L, G.G, A.M, P.P.L, M.M, A.S, S.B, A.B, V.F, W.L, J.X, and A.A contributed to bioinformatics analysis. J.F, B.W.L, J.N, C.L, M.L, S.L, and T.G generated figures. J.F, B.W.L, J.N, C.L, S.L, T.G, M.L, J.G, V.K, C.P, C.W, and J.X contributed to writing and editing the manuscript.

## Supplementary Methods

### Whole Methylome Sequencing Across Centers

#### Short-read sequencing details

The short-read sequencing libraries were collected from participating laboratories and sequenced centrally on NovaSeq 6000 systems at one or two sequencing centers.

Libraries were pooled by library type in high concentration equimolar stock pools (4 nM). After pooling, bead-based clean-up was performed to remove peaks <200 bp. Briefly, 0.7 X volume of NEBNext Sample Purification beads was added to the pools and incubated for 10 mins at room temperature. The beads were clarified by placing on a magnet and washed twice with freshly prepared 80% ethanol. Beads were allowed to dry for 2 mins and resuspended in 0.1 X TE. The cleaned stock pools were quantified on an Agilent Bioanalyzer using High sensitivity DNA chip.

#### Sequencing Center 1

Pooled libraries were diluted to 1.5 nM. were loaded on a NovaSeq S4 flowcell with a final loading concentration of 250 pM for all libraries with the exception of EM-Seq, which was loaded at 300 pM. Unrelated standard libraries were added at 5% instead of PhiX to balance the base composition during sequencing. All libraries were sequenced PE150 according to the manufacturer’s instructions (Illumina) with targeted per replicate CG coverage of 20x.

Base calling was performed using RTA v3.4.4 In cases where libraries were not prepared with dual-unique indices, they were demultiplexed using the expected index 2 sequence derived from the universal adapter. Demultiplexing and fastq generation was performed using Picard 2.20.6 using default settings except as listed below:

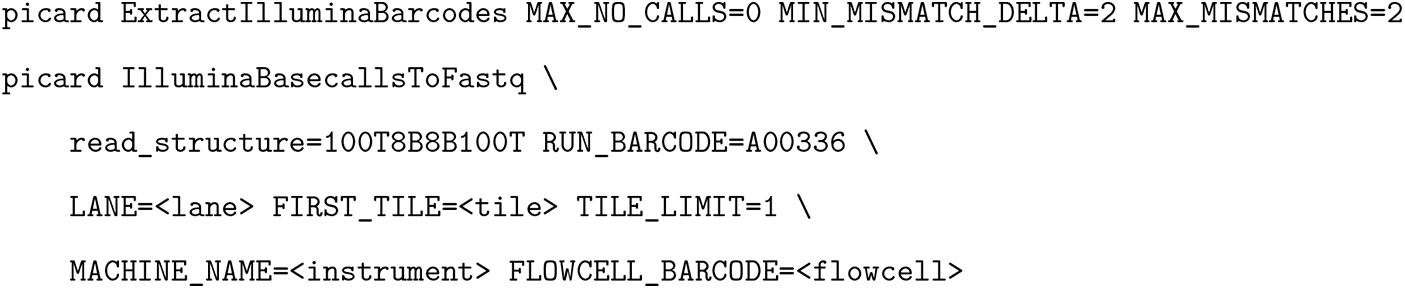

#### Sequencing Center 2

The high concentration equimolar stock library pools were sent to Illumina in order to ameliorate depth of sequencing for the WGBS libraries. Libraries pools were diluted to 1.5 nM and a final loading concentration of 300 pM was loaded on the flow cell with 5% PhiX. The libraries were sequenced on an Illumina NovaSeq 6000 S4 flowcell with direct flow cell loading (XP workflow) according to manufacturer’s instructions. MethylSeq, SPLAT and TruSeq pools were multiplexed on two lanes; SPLAT libraries on their own in the third lane; and TrueMethyl libraries on their own in the fourth lane. Base calling was performed using RTA v3.4.4. Run data were uploaded to BaseSpace and fastq files were generated using default parameters.

## Supplementary Results

### Alignment and Methylation Caller Comparisons

The first step after data QC was to map reads to a reference genome and estimate levels of methylation per CpG. We evaluated the performance of commonly used alignment/methylation calling packages, including Bismark [41], BitMapperBS [42], BSseker2 [43], bwa-meth [44], and gemBS [45]. For each software, we aligned reads to the GRCh38 human reference genome, with a set of bisulfite controls appended as additional contigs (see methods and Figure S2). We focused our analysis to Ashkenazi Son (HG002) data for these comparisons, using all replicates from each of the five short read epigenetic library types.

Although we successfully ran gemBS, its outputs were removed from further comparison for two reasons: (1) the maximum likelihood-based modeling of methylation percentages did not allow for merging of values across replicates, and (2) an unusually low percentage of CpGs were detected compared to all other platforms, prohibiting genome-wide comparison.

The mapping of reads showed aligner-specific distributions (Figure S3a). bwa-meth was able to map the highet percentage of reads to the reference genome, followed by bitmapperBS, BSSeeker2, and then Bismark. bwa-meth and Bismark tend to allow reads to align to multiple locations in the genome (marking these reads as secondary or supplementary alignments and ignoring them for methylation calling). BitMapperBS and BSseeker2 more commonly kept reads unmapped rather than align them ambiguously, although Bismark had the highest rate of unmapped reads. All four softwares had similar rates of duplicate read marking, except for BSseeker2 which tended to mark fewer reads as duplicates. It should be noted that an external program, Picard MarkDuplicates was used for deduplication in bwa-meth, BitMapperBS, and BSseeker2. Despite this, BSseeker2 samples still had fewer duplicate reads than other library types.

We then calculated the mapping effiency, defined as the percentage of bases aligned and retained for methylation calling (see below for the effects of read filtration) divided by the total bases per replicate (Figure S3b), as well as the mean coverage achieved per CpG dinucleotide (Figure S3c). bwa-meth returned both the most efficient mapping rate, as well as the highest mean coverage per CpG within every dataset except for TruSeq, where outputs from each software matched very closely. Generally, BitMapperBS scored second in efficiency and depth of coverage, followed by Bismark, then BSseeker2.

The running time of each aligner was tested using one million random paired-end reads from each replicate and run ten times, summarized in Supplementary Table 1. BitMapperBS was the fastest aligner, with an average of 550-650 read pairs processed per CPU core per second, with stable performance between replicates. Bismark and bwa-meth showed equal alignment speed (about 200 read pairs per CPU core per second). However, Bismark showed the most variability of timing between runs.

We then tested the distribution of CpGs called by each software (Figure S3d) to look for any aligner-specific biases. All four programs returned a nearly identical distribution of CpGs called throughout the genome. The highest genomic enrichment was detected at 5’UTRs, protomer regions, and exonic regions by all programs. Therefore, even though mapping efficiency and CpG depth was influenced by software, the genomic distribution of CpGs was reliably called by all softwares examined.

As a result of these comparisons, outputs from bwa-meth were used for all downstream analyses.

### 5-hydroxymethylcytosine Detection

Total 5-methylcytosine (5mC) and 5-hydroxymethylcytosine (5hmC) levels within each cell line examined in this study were measured by LC-MS/MS (Supplementary Table 6). The estimated percentage of 5hmC levels across all seven cell lines were below the limit of detection for this method.

In order to validate these results at base-level resolution, we used the NuGEN TrueMethyl oxBS-Seq library prepartion kit (aka TrueMethyl), which allows investigators to measure 5mC and 5hmC in an indirect manner on the sequence level. For completeness, each cell line replicate was processed using both bisulfite only (BS = 5mC + 5hmC) and an oxidative reaction prior to sodium bisulfite treatment (OX = 5mC).

Figure S12 shows that all cell lines have a higher level of 5mC compared to 5hmC (Figure S12a,b). The low 5hmC levels were also observed at the single-nucleotide resolution level, with similar correlations between the two library preparations across all cell lines (Figure S12c), and also within each cell lines (Figure S12d), where the PCA plot shows little to no separation between libraries prepared using BS or OX protocols.

As stated above, preparation of BS and OX libraries in parallel allows the determination of 5mC, 5hmC and C. We used the MLML2R package to estimate the level of each cytosine state, for each CpG sequenced, using HG002 as example (Figure S12e). The top panel shows that some CpG sites not only show 100% of a specific cytosine mark (C = 100% unmethylated CpG, mC = 100% methylated CpG), but also a mixture of two (mC_C = methylated or unmethylated C; hmC_C = hydroxymethylated or unmethylated C; mC_hmC = methylated or hydroxymethylated C) or of all cytosine mark (mC_hmC_C). Consistent with the LC-MS/MS quantitation, hmC marks were found in low proportions at some CpG sites. The results observed for HG002 were representative of all the 7 cell lines.

### Biological Significance of Between-Family Trio Differential Methylation

To determine the biological relevance of our results, we considered 51 CpGs on Chromosome 1 that had been previously identified as differentially methylated in an array analysis of approximately 300 individuals from Caucasian-American, African-American, and Han Chinese-American populations [46]. Annotation and methylation results from all 51 CpGs are available within Supplementary Table 5. Of the 7 sites with reported |PMD|>0.2 (Percent Methylation Difference) between Chinese-Americans and Caucasian-Americans, all had corresponding |PMD|>0.2 within the the microarray data. Additionally, 4 of these were identified as statistically significant DMAs across all six sequencing assays (five short read library types and Oxford Nanopore). Of the three remaining sites, the first (on the TAS1R3 promoter) was significantly hypomethylated in the Chinese family for EMSeq, Nanopore, SPLAT, and TrueMethyl, the second (on the PM20D1 promoter) had insuficient read coverage for TruSeq but was a DMA for the remaining assays, and the third (located on the C1orf100 promoter) was identified as a DMA for only SPLAT although estimated PMD values were greater than 0.1 for all assays. Notably, these sites were identified as methylation quantitative trait loci (meQTL) in the original analysis. In addition to TAS1R3, which is a sweetness taste receptor that is known to vary phenotypically between the Asian and Caucasian populations [47], there was strong concordance for 6 CpGs on the PM20D1 promoter, a gene associated with obesity and Alzheimer’s disease with demonstrated population-based variation [48, 49].

We additionally reviewed the collection of 29,802 sites on Chromosome 1 that were identified as differentially methylated for four or more of the six sequencing assays. Following annotation with HOMER [50], analysis with DAVID [51] identified a subset of 133 genes associated with hypertension (Benjamini-Hochberg adjusted *p*-value = 5.0E-13), 54 genes associated with osteoporosis (*p* = 5.0E-13), and 18 genes associated with atopic dermatitis (*p* =1.0E-5) according to the GAD database [52]. Only 1204 (4.0%) of these sites were included on the Infinium MethylEPIC array, and while annotation for these sites included 53 of the hypertension-associated genes (*p*=3.3E-4) and 9 of those associated with atopic dermatitis (*p*=0.03), only 17 of the genes identified with osteoporosis were included and this was an insufficient number to result in a significant association.

### EMSeq Input Titration

In order to investigate the impact of input DNA on detection and characterization of CpG methylation, we generated EM-Seq libraries using 10ng, 50ng, and 100ng aliquots of input DNA for each replicate for each member of the Chinese Han Trio in this study (HG005-7). We then randomly subsampled each run *in silico* to a random set of 1M, 5M, 10M, 25M, 50M, and 100M paired end 150bp reads per input. At the lowest read input, the less complex 10ng library covered CpGs greater than 50ng and 100ng libraries, though beyond 25M paired end reads the more complex (50/100ng) libraries surpassed the 10ng library in mean CpG coverage (Figure S13a). All three library types exhibited similar distributions of CpG coverage across read titrations, reflecting fringe technical noise contributing to mean depth differences at low inputs that were evened out with more input. This was further validated by looking at the intersection of CpGs covered by each input type at each read filtration titer, where by 10M paired end reads the majority of sites were shared by all libraries, and notably the lowest input consistently covered the fewest unique CpGs (Figure S13c).

### Methyl EPIC Capture Correlations

We compared the whole epigenome libraries to sequencing replicates of Illumina Methyl Capture EPIC, a reduced representation bisulfite approach interrogating roughly 3.3 million CpGs with a preference for CpG islands and promoter regions. Results shown for HG002 are representative of all seven genomes. Methylation percentage of CpGs within replicates of Capture EPIC were compared to shared sites among whole methylome assays as well as Nanopore sequencing, with good Pearson correlation for all comparisons (average r=0.85). Capture EPIC tended to overestimate fully methylated sites that were estimated to be closer to 50-90% in other assays (Figure S14a).

Using 20X downsampled methylation data, the shared CpG coverage on Chromosome 1 in Capture EPIC sites was highly consistent with overall methylome coverage (Figure 2). Nanopore missed the fewest sites covered by EPIC (n=5,179), while TruSeq missed the most (n=21,712).

## Supplementary Figures

**Figure S1:**
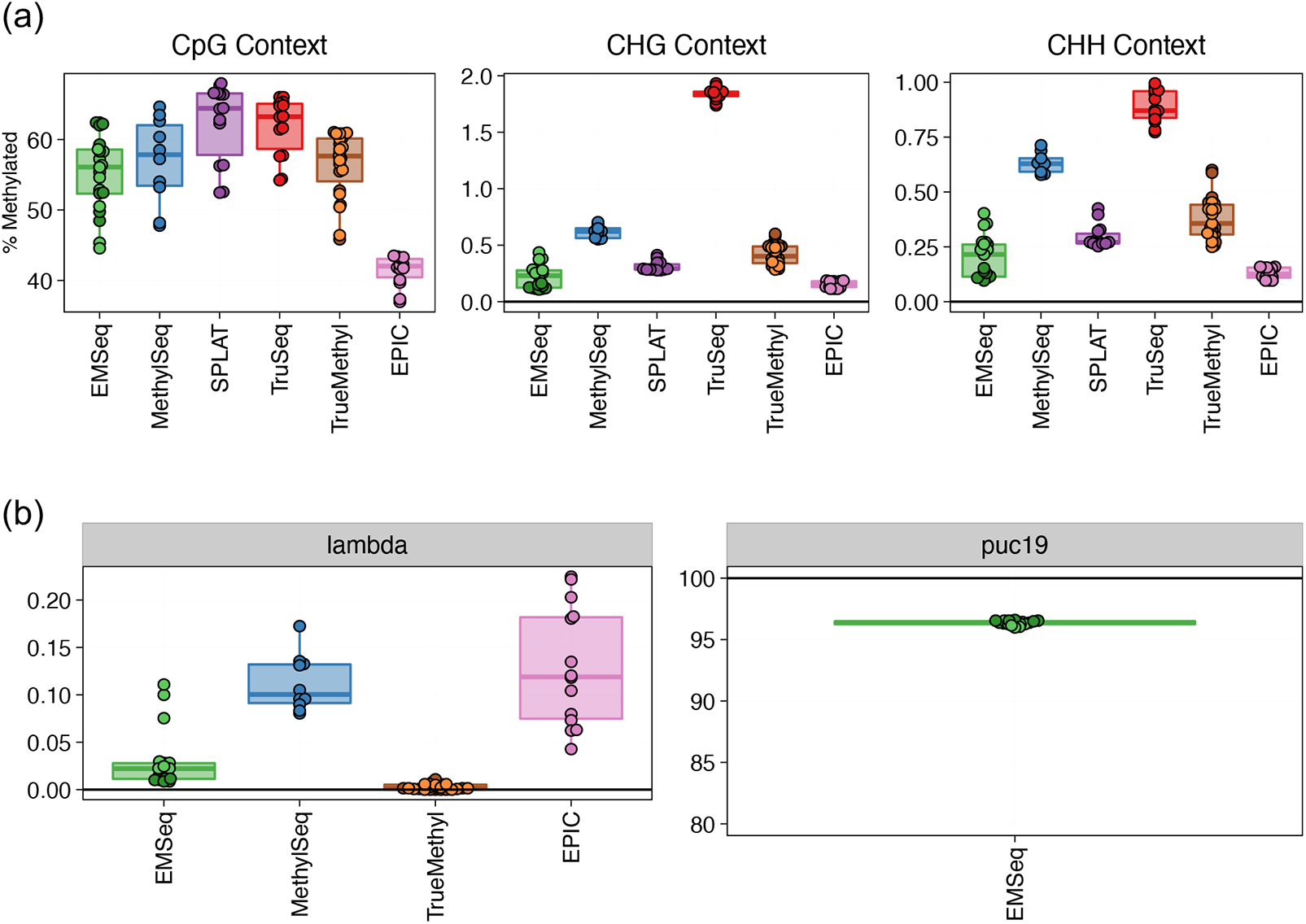
Measurement of sequencing control samples (a) Estimated methylation percentage in CpG, CHG, and CHH contexts per assay. Efficient conversion results in near-zero converted cytosines in CHG and CHH contexts. (b) Estimated methylation percentage in unmethylated controls, showing only assays that had these controls spiked in as a part of their library preparation.

**Figure S2:**
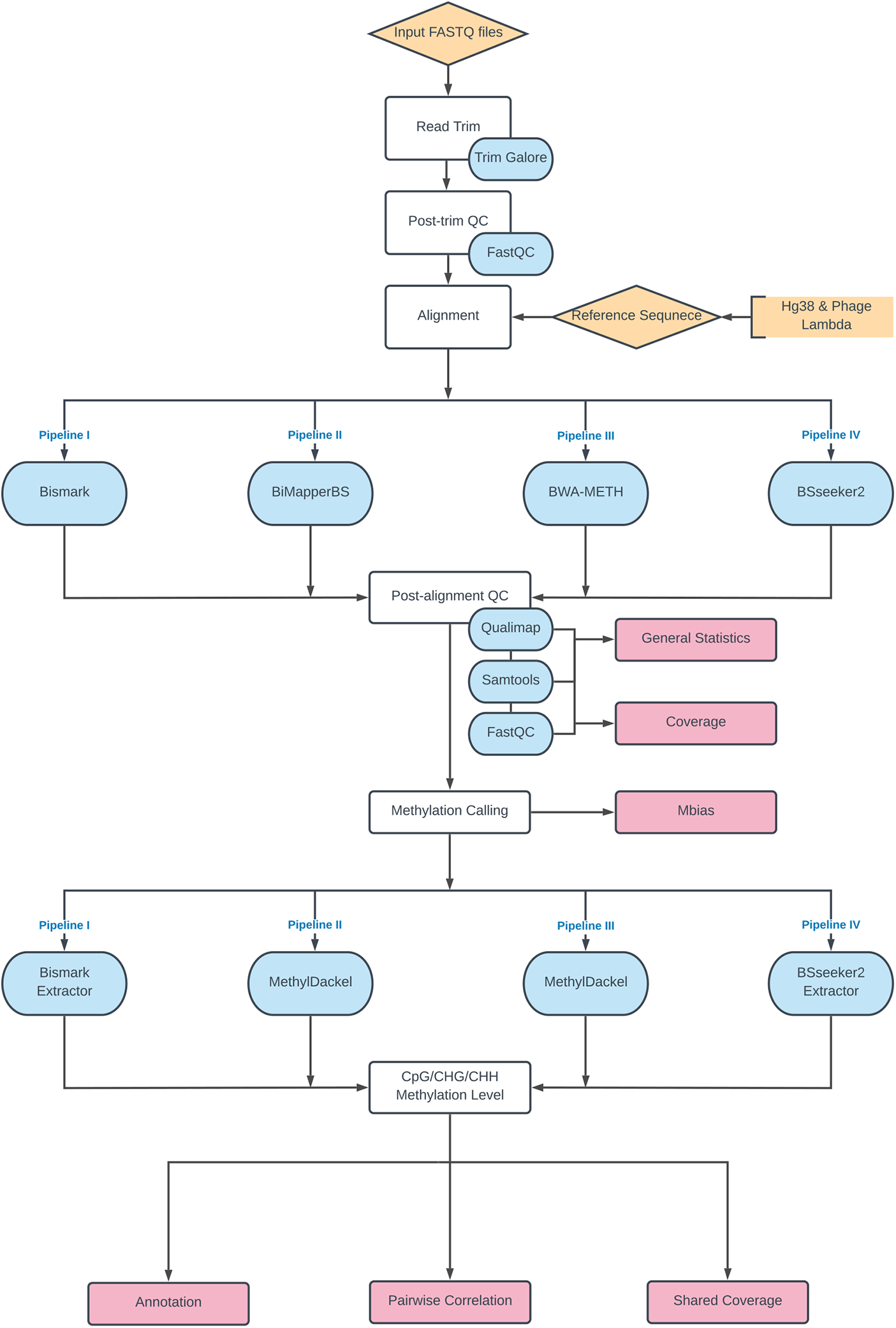
Flowchart showing recommended steps for read quality control, reference-based read alignment, and methylation extraction, for each methylation package analyzed.

**Figure S3:**
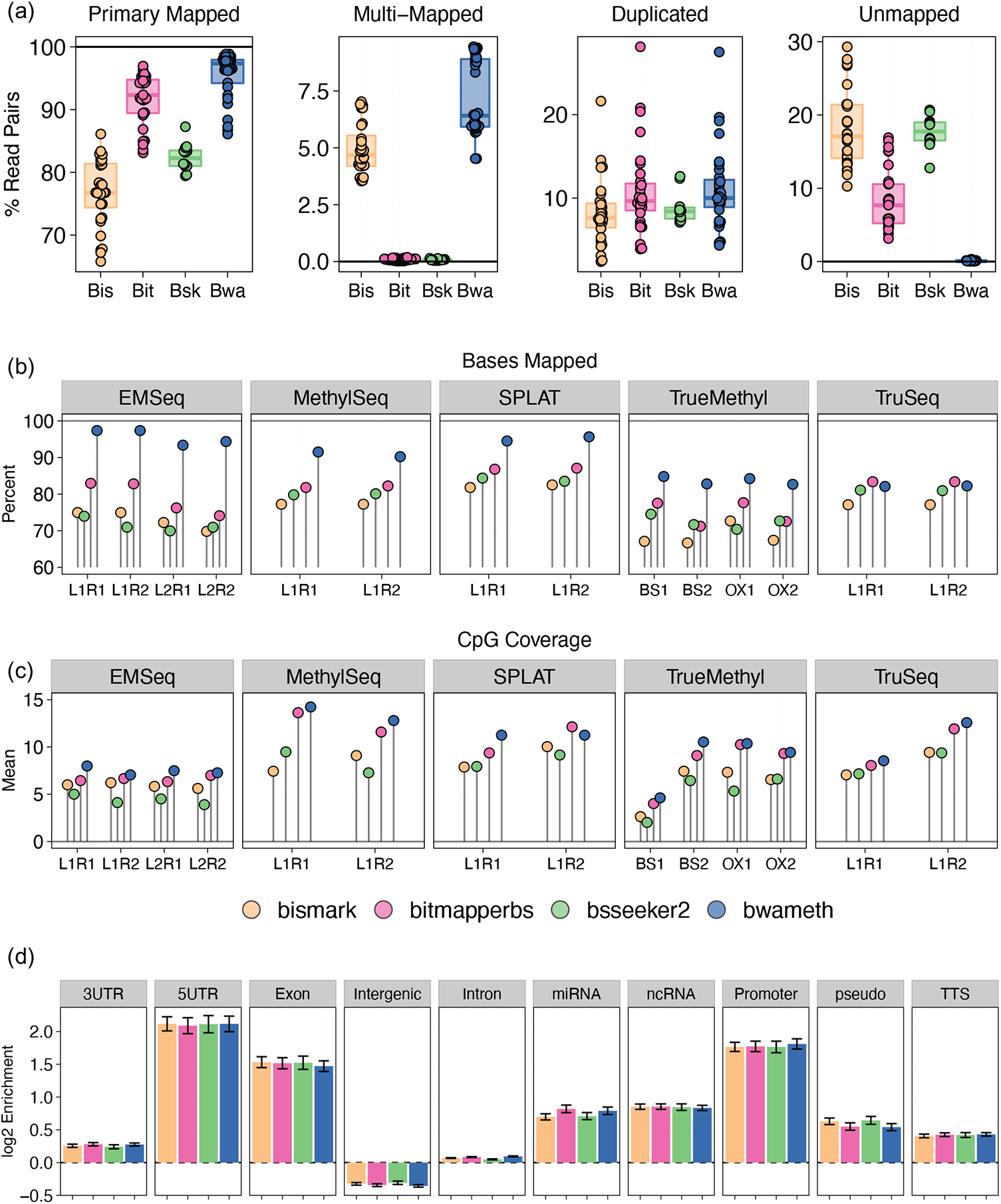
Comparison of outputs for each methylation detection pipeline. All figures show analysis of all HG002 samples for each short read epigenomic assay. (a) Distribution of reference-based read alignment outcomes, including primary mapped reads (both mates mapped in correct orientation within a certain distance), multi-mapped reads (read pairs containing secondary or supplementary alignments), reads marked as PCR or optical duplicates, and unmapped reads. Ambiguous and duplicate reads can be a subset of properly aligned reads. (b) Mapping efficiency per pipeline as measured by the total percentage of reads aligned to the reference genome. L1 and L2 = Lab 1/2; R1 and R2 = Replicate 1/2; BS1 and BS2 = bisulfite treatment replicates 1/2; OX1 and OX2 = oxidative-bisulfite replicates 1/2. (c) The mean coverage per CpG across the genome per pipeline. (d) The regions of the genomes covered per pipeline, measured as log2 enrichment against a null genomic distribution.

**Figure S4:**
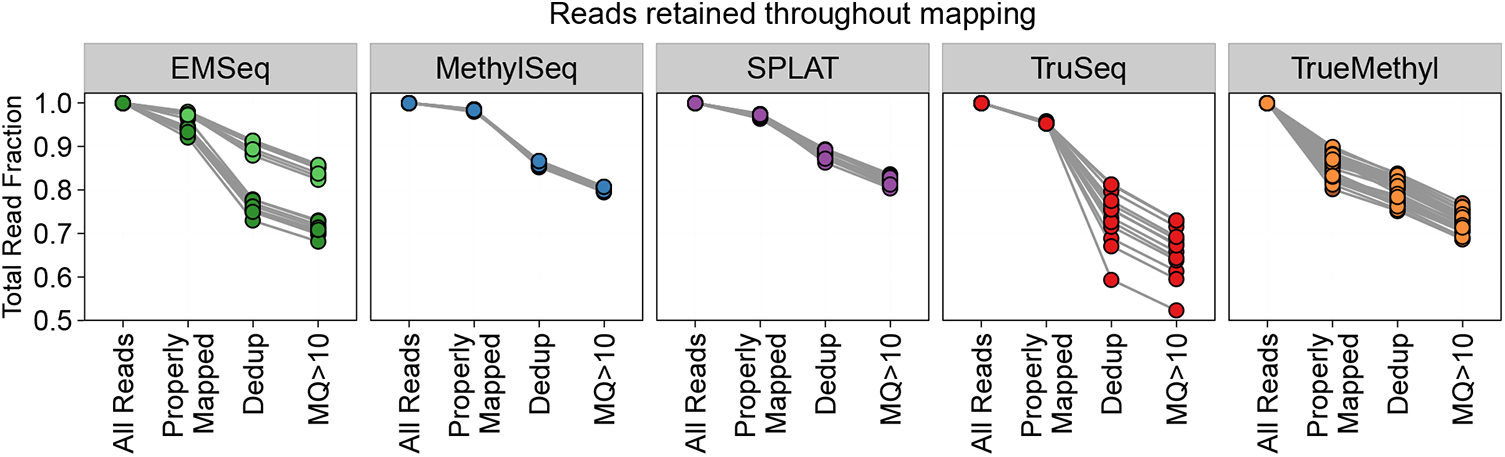
Read retention rate. The fraction of total reads that are retained after each step of the epigenome alignment process is shown per assay. Properly mapped = both mates of a pair were mapped in the correct orientation within a 1kb distance. Dedup = removing reads that are marked as duplicates. MQ = Mapping Quality.

**Figure S5:**
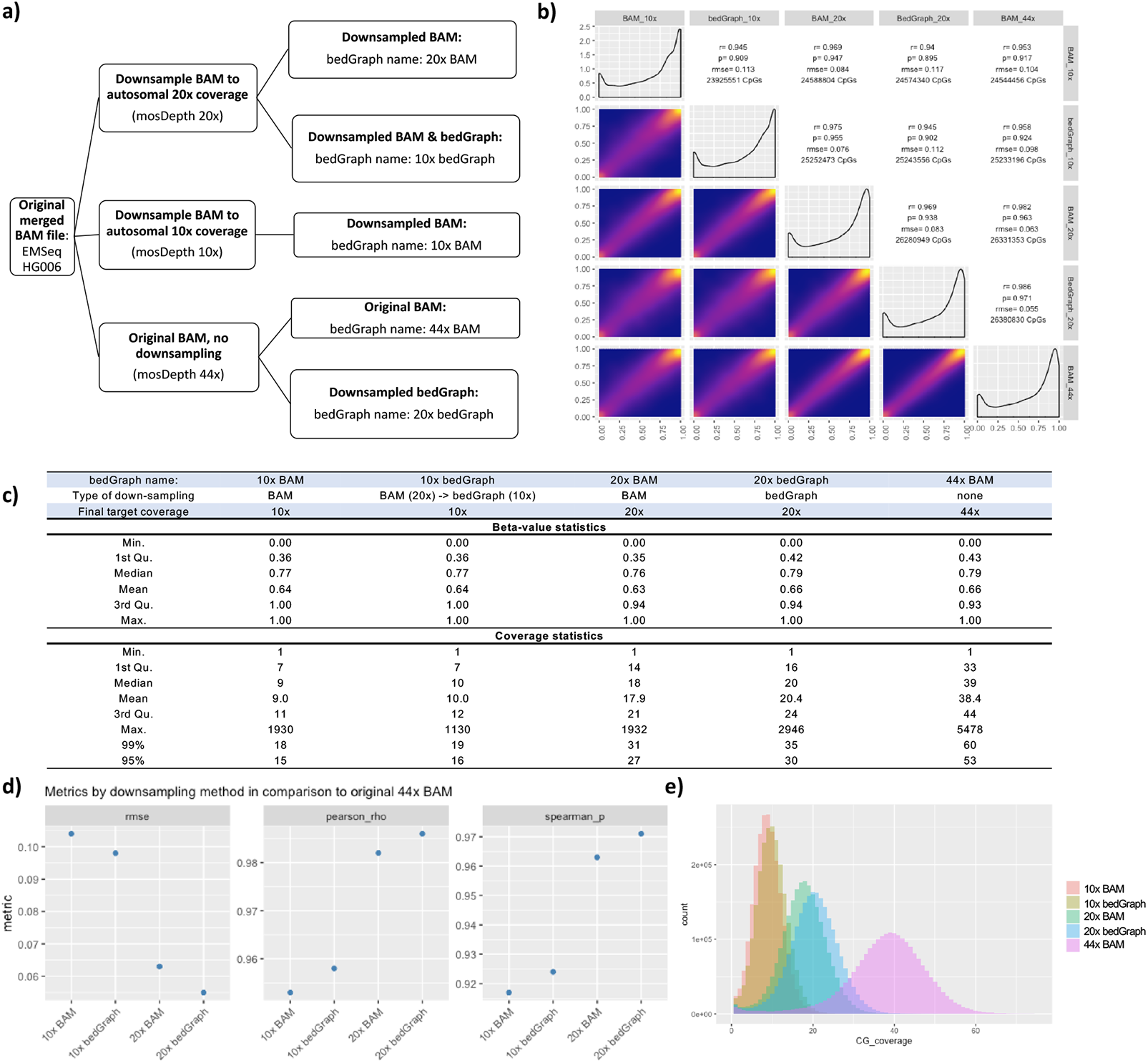
Downsampling evaluation for EMSeq / HG006. (a) Outline of the downsampling procedure and naming scheme of the downsampled libraries. (b) Pairwise correlation matrix of methylation values for the EMSeq HG006 library from Lab 1. Scatter plots of the methylation values are shown in the lower left. Histograms of the methylation values per library are shown across the diagonal. Pairwise Pearson (rho) and Spearman (p) correlation coefficients, root mean square error (RMSE), and the number of CpG dinucleotides with >= 5x coverage in both libraries are shown in the upper right. (c) Statistics over the methylation percentage distributions and observed read coverage of CpG sites in the various bedGraph files. (d) RMSE, Pairwise Pearson (p) and Spearman (rho) correlations between downsampled BAM and bedGraph files in comparison to the original 44x average coverage BAM file. (e) Histograms of the CG dinucelotide read coverage of each bedGraph file prior (44x BAM) to and after downsampling the BAM or bedGraph.

**Figure S6:**
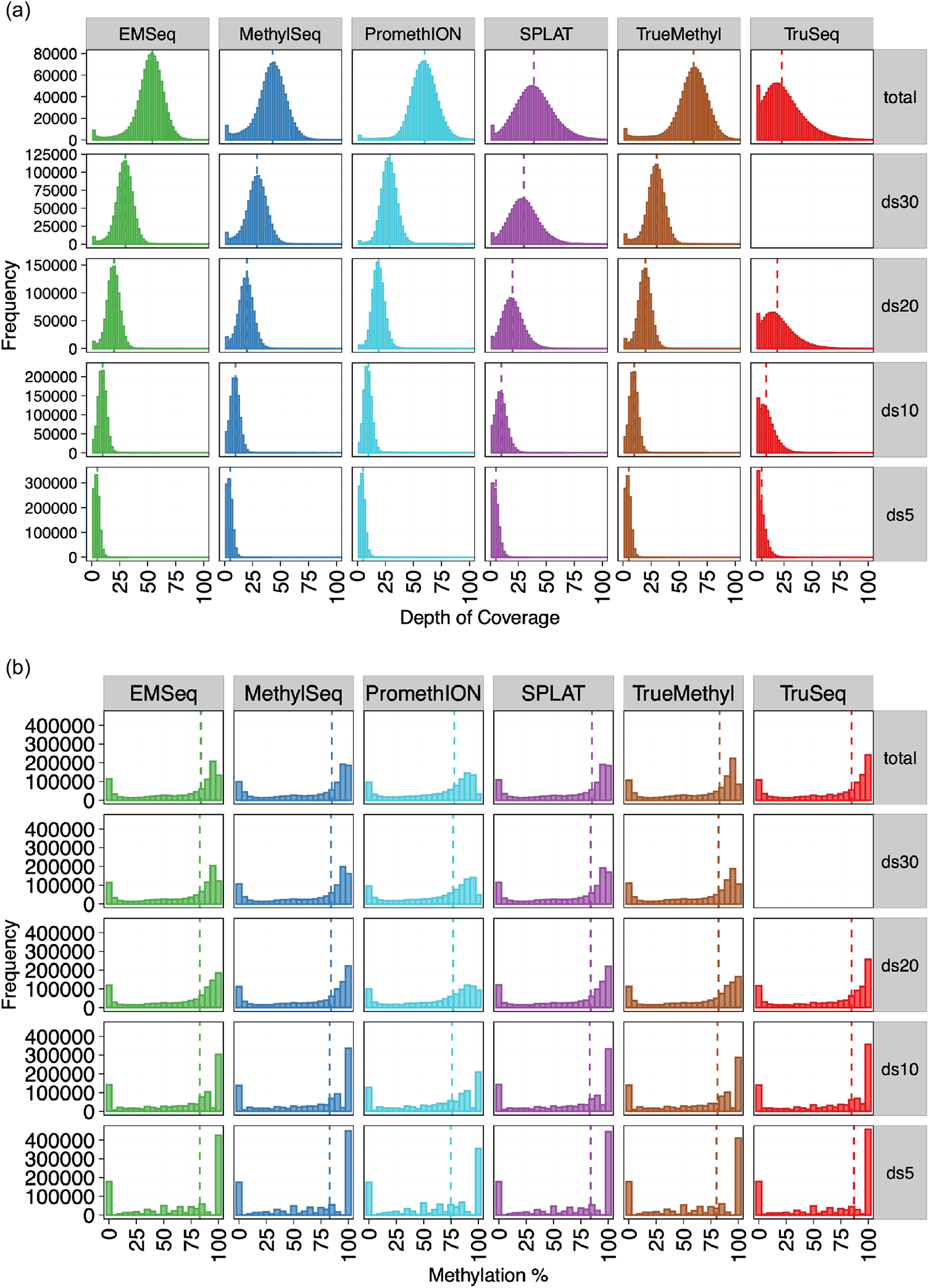
CpG coverage and methylation percentage distributions for complete and downsampled libraries per assay. All values are shown for replicates of HG002. ds = downsample, indicating the mean CpG coverage samples were normalized to. Vertical dotted lines indicate median coverage/methylation percentage.

**Figure S7:**
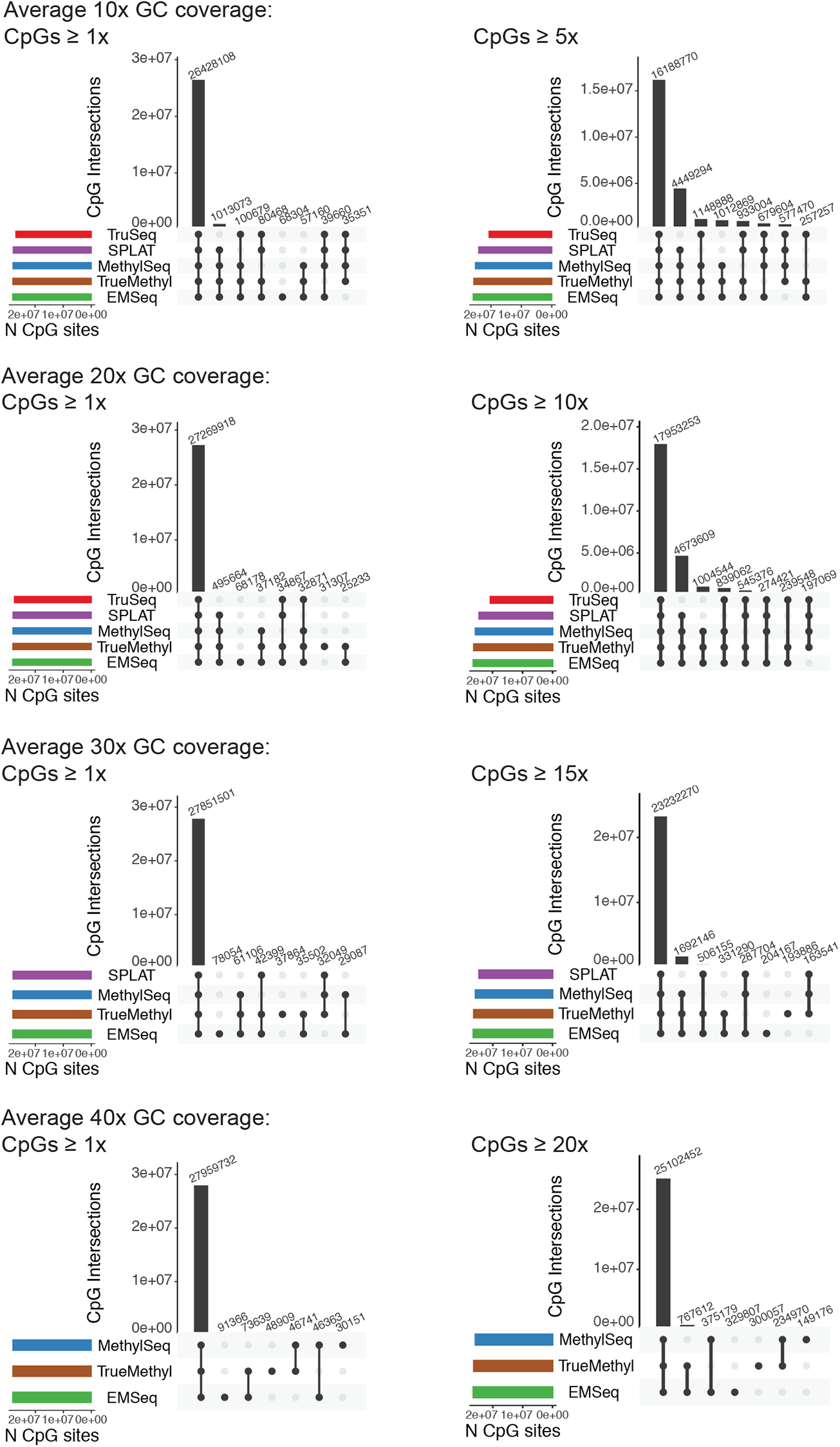
UpSet plots showing shared coverage of CpGs across assays across downsampling schema, with a minimum of 1x cov per CpG on the left and a minimum of 50% of the downsampling scheme on the right (e.g. minimum of 5x coveage for 10x downsampled data).

**Figure S8:**
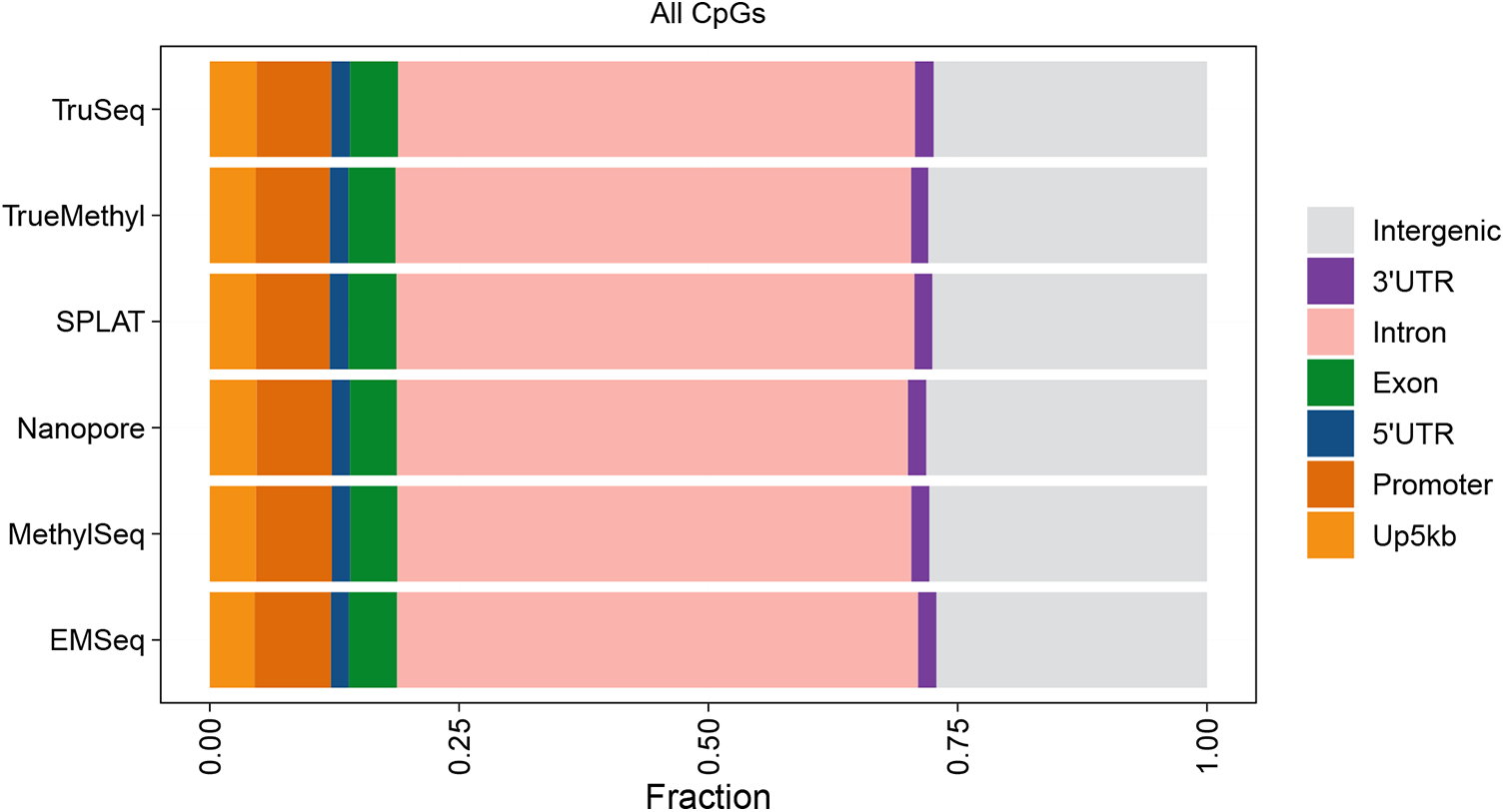
Annotating CpGs covered by each assay using normalized mean 20x coverage data, showing the consistency of coverage genome-wide. Up5kb = 5kb upstream of promoter regions. Promoter = 1kb upstream of transcript start sites.

**Figure S9:**
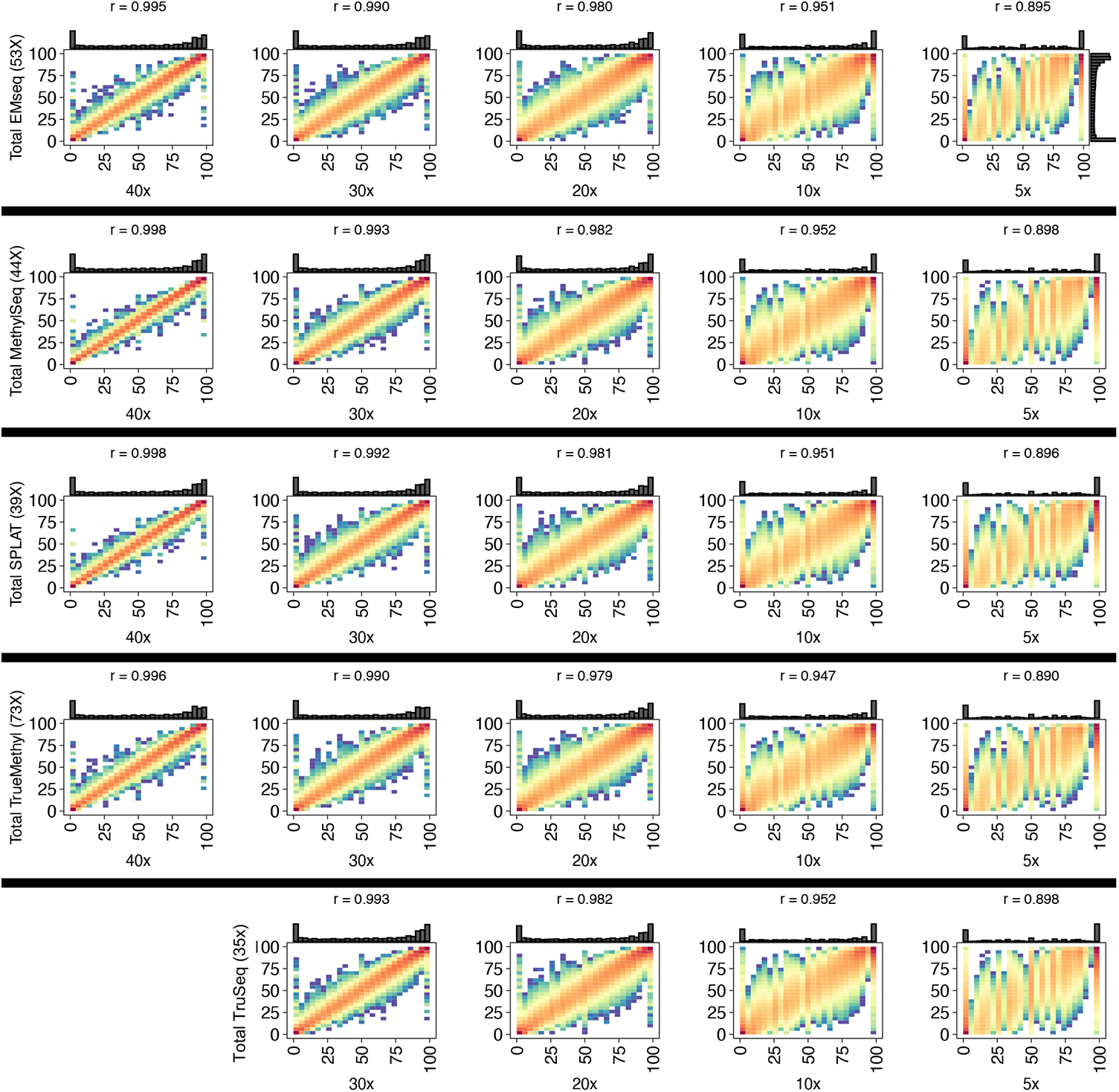
Pearson correlations of methylation percentage estimation within each assay, comparing the total data (y-axes) against their respective downsampled schema (x-axes), for combined replicates of HG002 libraries. Pearson values are shown above each comparison, as well as marginal histograms showing methylation percentage distributions. For TruSeq, the total data returned a mean coverage of 35X, meaning that a comparison to 40X downsampling was not possible.

**Figure S10:**
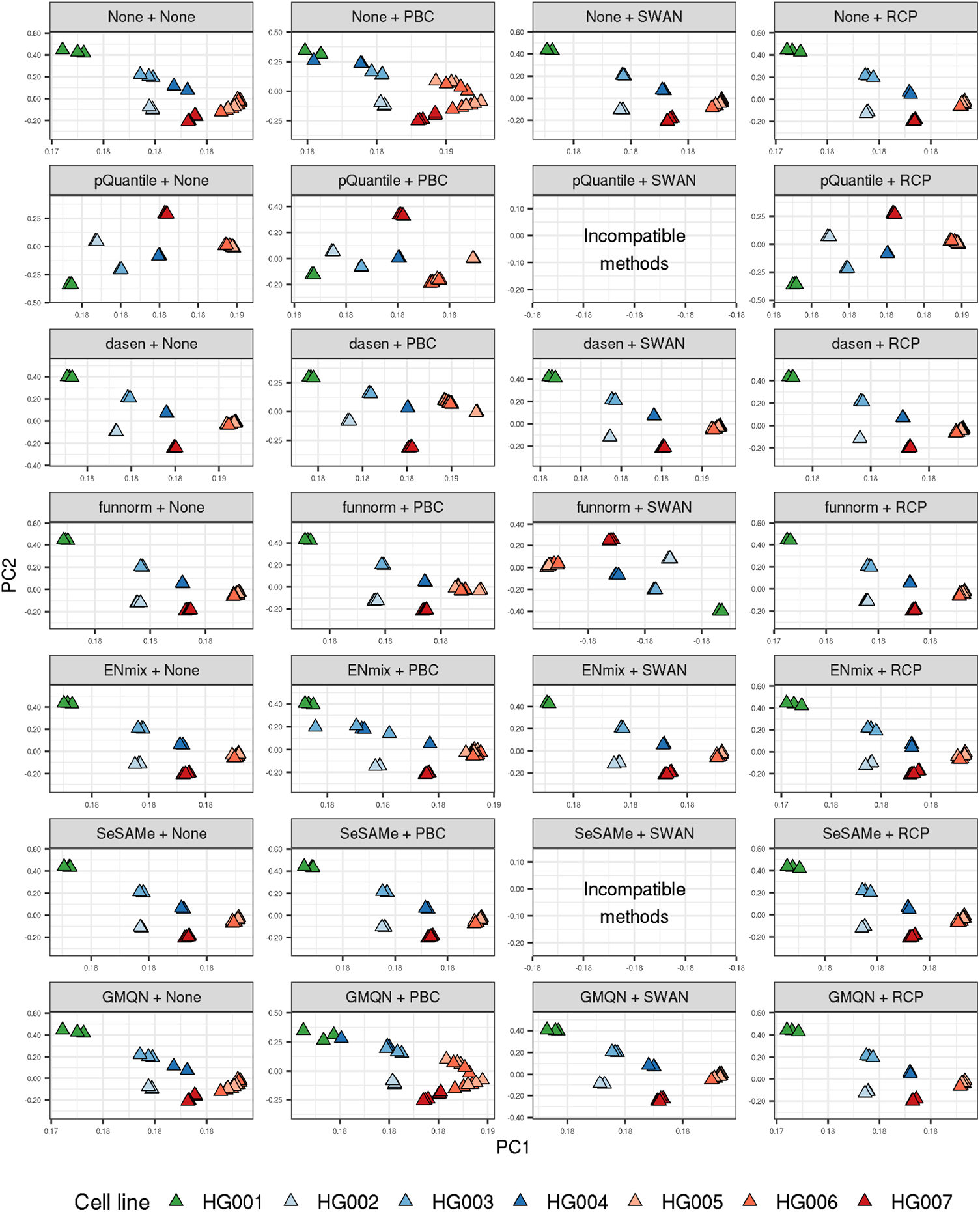
First two principal components (PCs) calculated from 678,597 CpG sites with complete information in all normalized microarray datasets, by normalization pipeline.

**Figure S11:**
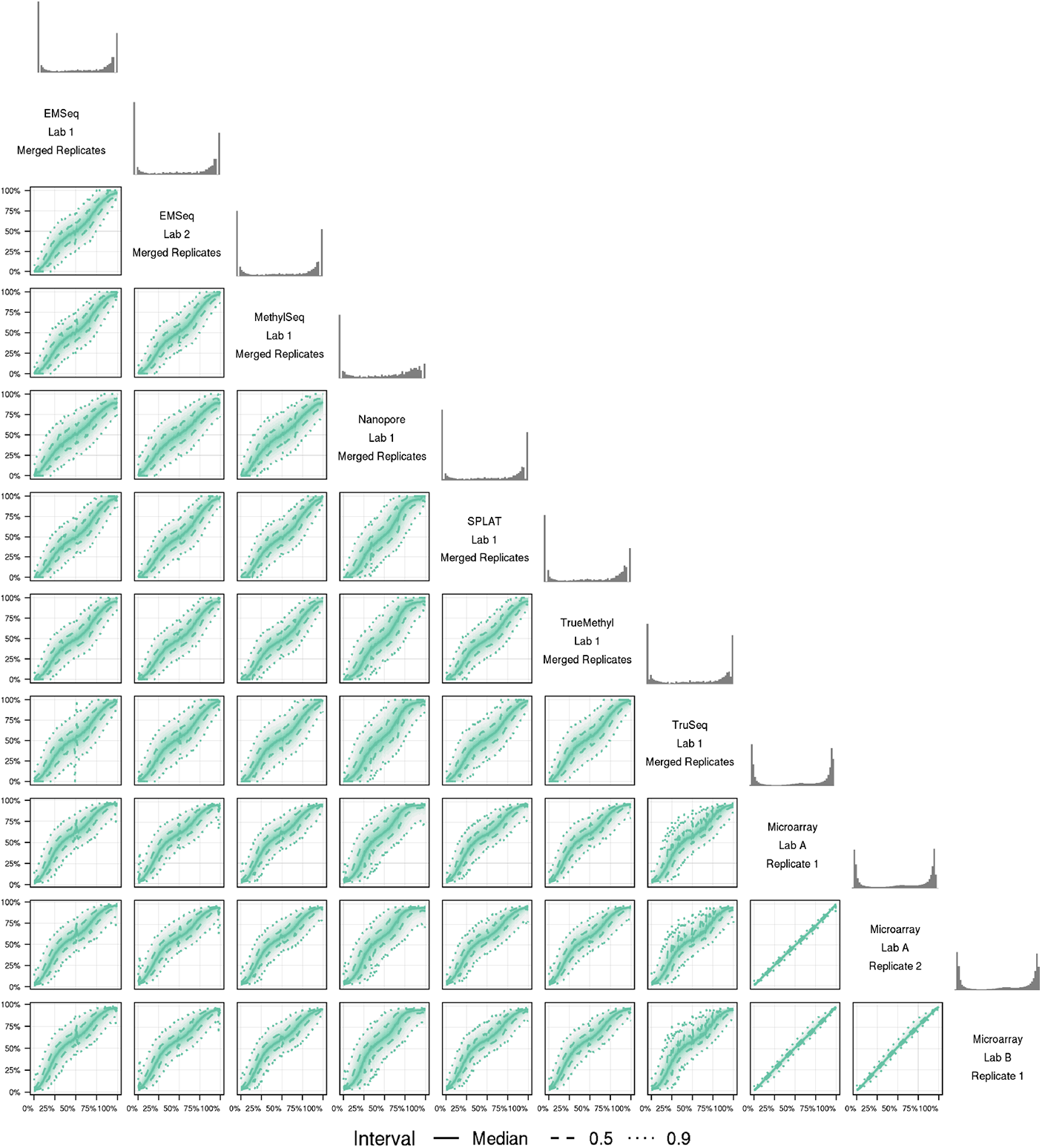
Distribution of beta values across HG002 samples at 841,833 CpG sites with complete information in all assays. Beta values for the assay on the x axis were binned (binwidth=0.01) to calculate beta value deciles for the assay on the y axis, indicated by the color transparency. 90% of the y-axis values fall between the outermost dotted lines for each bin along the x-axis. Marginal histograms for each assay are shown above the assay label.

**Figure S12:**
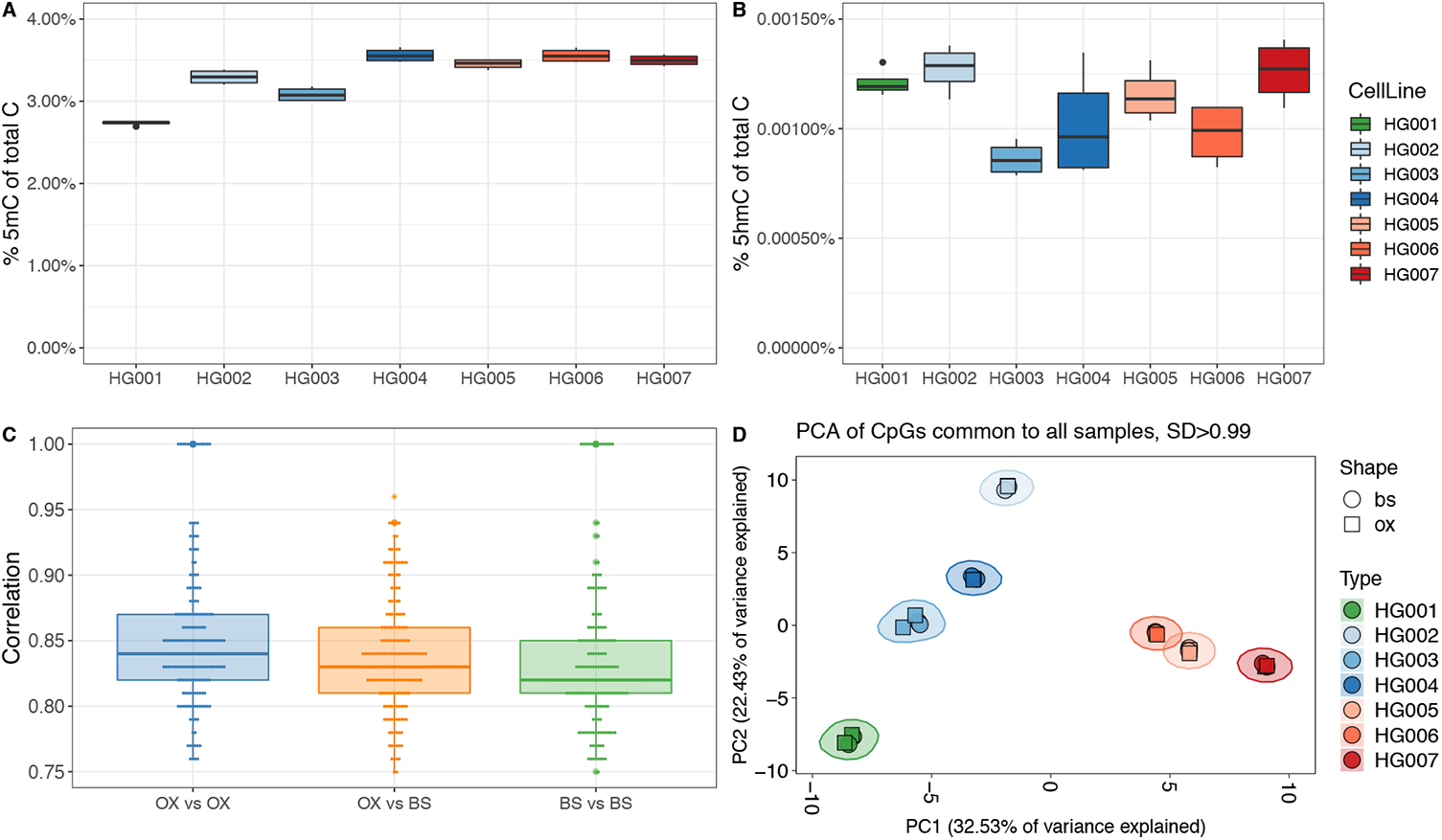
Capture of 5mC and 5hmC from TrueMethyl replicates, including bisulfite-only (bs) and oxidative bisulfite (ox). (a) Percent of inferred 5mC among all cytosines in the genome. (b) Percent of inferred 5hmC among all cytosines in the genome. (c) Pearson correlation of replicates across genomes between oxidative and bisulfite replicates. (d) Unsupervised clustering of samples, including OX and BS samples.

**Figure S13:**
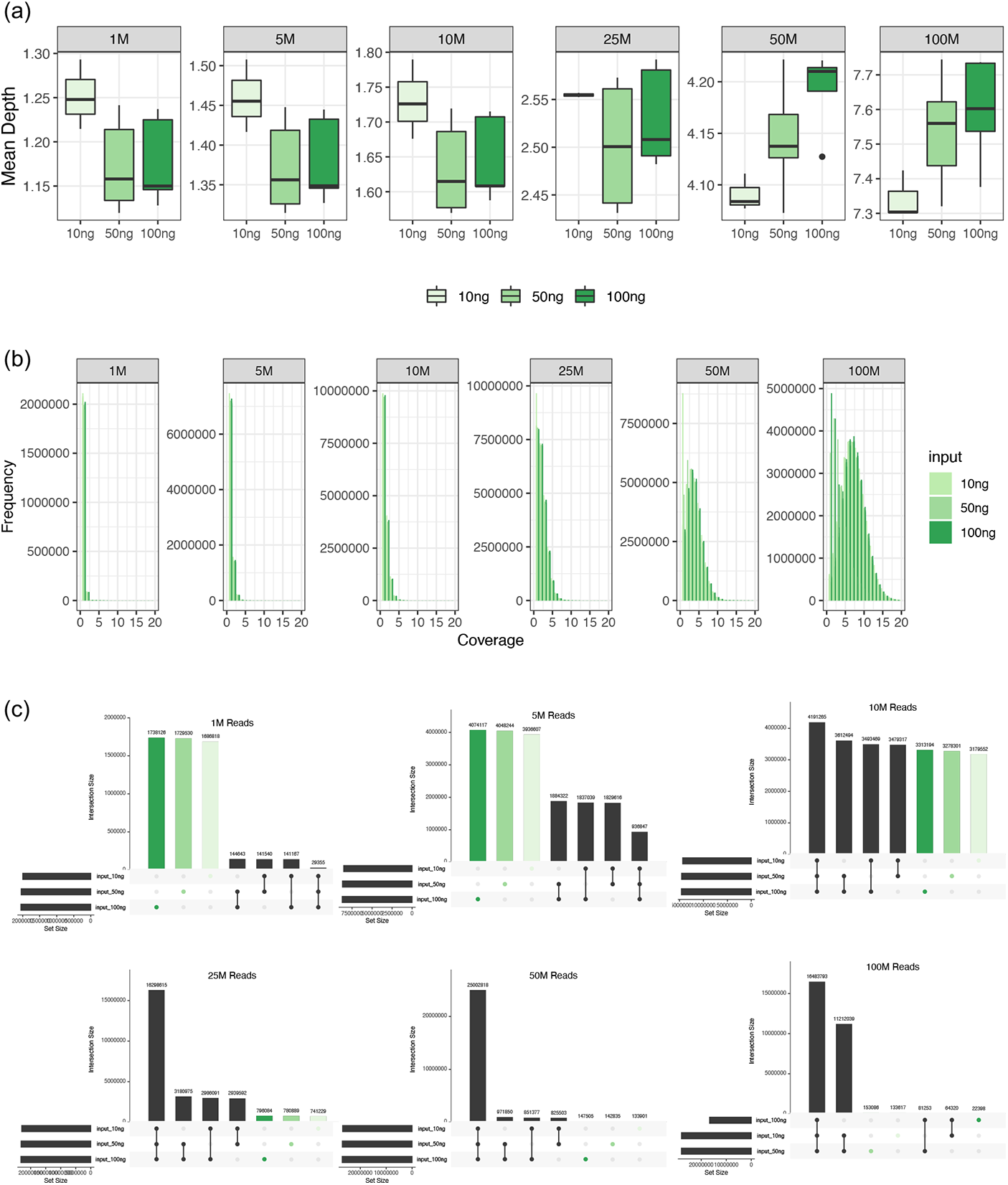
EM-Seq read titration experiment. Replicates generated using 10ng, 50ng, and 100ng of input DNA for HG005, HG006, and HG007 were randomly downsampled to 1M, 5M, 10M, 25M, 50M, and 100M paired end 150bp input reads. (a) Distribution of mean depth of CpGs covered for each input amount. (b) Read coverage distributions per input type per downsampled read amount. (c) UpSet plots showing the intersections of CpGs shared by each downsampling scheme, as well as uniquely covered CpGs.

**Figure S14:**
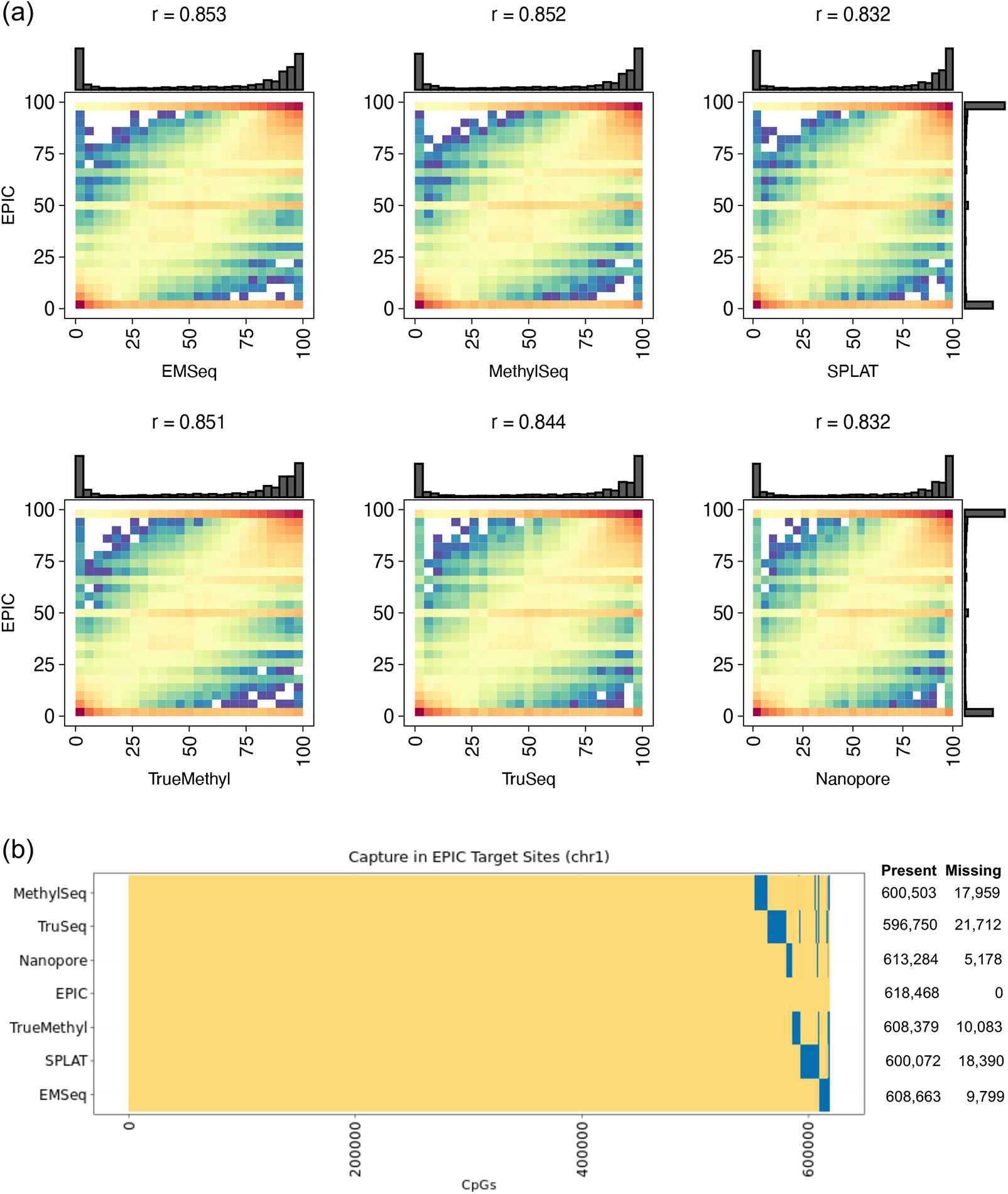
(a) Pearson correlation of percent methylation estimates of Methyl Seq EPIC Capture versus each whole methylome library. All values are shown for Chromosome 1 of HG002 replicates. (b) Distribution of CpGs covered (in yellow) or missed (in blue) by each assay on Chromosome 1. Total values are shown per assay in the table on the right.

## Supplementary Tables

**Supplementary Table 1.** Timing comparison of each alignment and methylation estimation software. Ten permutations of each software were run, and the mean times are reported, as well as the standard deviation per sotware.

| Step | Software | No. Reads | Mean Time (min) | Standard Dev (min) |
| --- | --- | --- | --- | --- |
| Alignment | Bismark | 1M | 19.33 | 1.74 |
|  | BitmapperBS | 1M | 11.98 | 3.76 |
|  | BSseeker2 | 1M | 65.76 | 3.5 |
|  | bwa-meth | 1M | 29.92 | 6.17 |
| Meth Calling | Bismark | 1M | 2.86 | 0.9 |
|  | BitmapperBS | 1M | 1.97 | 0.17 |
|  | BSseeker2 | 1M | 39.87 | 17.4 |
|  | bwa-meth | 1M | 0.24 | 0.07 |

**Supplementary Table 2.** Read and mapping statistics for all cell lines. stDev = standard deviation. Values shown are averages across replicates for each library. Useable bases are calculated as the total mapped bases as a percentage of the total number of bases sequenced.

| **Insert Size** | **HG001** | **HG002** | **HG003** | **HG004** | **HG005** | **HG006** | **HG007** | **mean** | **stDev** |
| --- | --- | --- | --- | --- | --- | --- | --- | --- | --- |
| EMSeqLAB01 | 297.1 | 299.6 | 349.75 | 288.95 | 300.53 | 298.23 | 330.3 | 309.21 | 22.11 |
| EMSeqLAB02 | 325.8 | 327.95 | 326.35 | 328.15 | 320.2 | 325.47 | 358.57 | 330.35 | 12.72 |
| MethylSeq | 249.77 | 250.93 | 250.88 | 247.13 | 249.67 | 256.45 | 247.78 | 250.37 | 3.04 |
| SPLAT | 214.83 | 217.81 | 220.71 | 223.74 | 220.34 | 223.85 | 224.18 | 220.78 | 3.52 |
| TruSeq | 218.58 | 217.68 | 219.88 | 216.77 | 214.67 | 217.58 | 215.78 | 217.28 | 1.74 |
| TrueMethylOX | 207.7 | 203.83 | 209.95 | 205.93 | 211.8 | 211.5 | 204.4 | 207.87 | 3.3 |
| TrueMethylBS | 228.6 | 220.67 | 226.5 | 224.2 | 224.73 | 227.4 | 219 | 224.44 | 3.52 |
| EPIC | 225.7 | 227.45 | 225.65 | 227 | 229.65 | 229.7 | 227.7 | 227.55 | 1.65 |
| **Primary Mapping %** | **HG001** | **HG002** | **HG003** | **HG004** | **HG005** | **HG006** | **HG007** | **mean** | **stDev** |
| EMSeqLAB01 | 97.48 | 97.63 | 97.37 | 97.87 | 97.14 | 97.36 | 97.32 | 97.45 | 0.24 |
| EMSeqLAB02 | 94.89 | 93.27 | 94.09 | 94.46 | 91.97 | 94.27 | 93.33 | 93.75 | 0.98 |
| MethylSeq | 98.4 | 98.36 | 98.32 | 98.41 | 98.16 | 98.36 | 98.39 | 98.34 | 0.09 |
| SPLAT | 96.44 | 97.18 | 96.92 | 97.21 | 97.07 | 97.48 | 97.24 | 97.08 | 0.33 |
| TruSeq | 95.33 | 95.45 | 95.23 | 95.58 | 95.51 | 95.55 | 95.37 | 95.43 | 0.13 |
| TrueMethylOX | 85.95 | 84.53 | 87.43 | 87.01 | 85.64 | 87.15 | 85.11 | 86.12 | 1.11 |
| TrueMethylBS | 85.17 | 83.5 | 86.66 | 86.06 | 84.73 | 86.4 | 84.2 | 85.25 | 1.18 |
| EPIC | 98.27 | 98.31 | 98.37 | 98.12 | 98.28 | 98.27 | 98.23 | 98.26 | 0.08 |
| **Duplicate %** | **HG001** | **HG002** | **HG003** | **HG004** | **HG005** | **HG006** | **HG007** | **mean** | **stDev** |
| EMSeqLAB01 | 9.28 | 9.13 | 9.15 | 8.65 | 11.28 | 12.1 | 10.6 | 10.03 | 1.3 |
| EMSeqLAB02 | 23.61 | 25.03 | 23.97 | 23.37 | 27.08 | 23.68 | 25.11 | 24.55 | 1.31 |
| MethylSeq | 13.75 | 13.77 | 13.84 | 14.42 | 13.56 | 14.44 | 13.32 | 13.87 | 0.42 |
| SPLAT | 12.37 | 12.02 | 11.63 | 10.9 | 11.88 | 10.86 | 13.28 | 11.85 | 0.84 |
| TruSeq | 21.73 | 22.53 | 27.26 | 24.35 | 25.86 | 31.57 | 25.77 | 25.58 | 3.28 |
| TrueMethylOX | 21.24 | 21.66 | 18.19 | 19.48 | 20.86 | 19.56 | 21.21 | 20.31 | 1.26 |
| TrueMethylBS | 21.29 | 21.95 | 17.89 | 18.72 | 20.15 | 18.66 | 20.22 | 19.84 | 1.48 |
| EPIC | 66.42 | 67.43 | 62.9 | 63.92 | 69.21 | 69.5 | 68.24 | 66.8 | 2.56 |
| **Dinucleotide Bias** | **HG001** | **HG002** | **HG003** | **HG004** | **HG005** | **HG006** | **HG007** | **mean** | **stDev** |
| EMSeqLAB01 | 3.77 | 2.97 | 3.36 | 2.98 | 2.7 | 2.69 | 2.99 | 3.07 | 0.38 |
| EMSeqLAB02 | 1.12 | 0.84 | 1.16 | 0.98 | 1.14 | 1.11 | 1.1 | 1.06 | 0.11 |
| EPIC | 26.58 | 26.74 | 26.59 | 26.69 | 26.43 | 26.36 | 26.93 | 26.62 | 0.19 |
| MethylSeq | 3.4 | 3.71 | 3.43 | 3.41 | 3.47 | 3.53 | 3.52 | 3.5 | 0.11 |
| SPLAT | 6.95 | 7.58 | 5.55 | 6.36 | 6.66 | 6.24 | 5.8 | 6.45 | 0.69 |
| TrueMethylBS | 3.43 | 3.44 | 3.8 | 3.3 | 3.73 | 3.61 | 3.99 | 3.61 | 0.24 |
| TrueMethylOX | 3.3 | 3.71 | 3.64 | 3.71 | 3.72 | 3.72 | 3.71 | 3.64 | 0.16 |
| TruSeq | 24.66 | 23.09 | 24.1 | 23.69 | 23.13 | 23.36 | 23.83 | 23.69 | 0.56 |
| **Useable Bases %** | **HG001** | **HG002** | **HG003** | **HG004** | **HG005** | **HG006** | **HG007** | **mean** | **stDev** |
| EMSeqLAB01 | 90.21 | 90.35 | 90.26 | 90.71 | 87.88 | 87.61 | 88.89 | 89.42 | 1.27 |
| EMSeqLAB02 | 76.01 | 74.6 | 75.51 | 76.16 | 72.62 | 76.01 | 51.56 | 71.78 | 9 |
| EPIC | 33.2 | 32.21 | 36.69 | 35.67 | 30.46 | 30.17 | 31.4 | 32.83 | 2.52 |
| MethylSeq | 74.46 | 74.5 | 74.55 | 73.8 | 74.77 | 74.01 | 74.9 | 74.43 | 0.39 |
| SPLAT | 78.9 | 80.62 | 81.19 | 83 | 81.43 | 82.79 | 81.4 | 81.33 | 1.38 |
| TrueMethylBS | 70.21 | 66.6 | 71.08 | 70.13 | 70.03 | 71.69 | 71.97 | 70.24 | 1.78 |
| TrueMethylOX | 67.53 | 66.49 | 68.45 | 68.5 | 68.07 | 68.66 | 66.78 | 67.78 | 0.87 |
| TruSeq | 62.68 | 62.98 | 60.23 | 61.19 | 60.29 | 62.58 | 51.03 | 60.14 | 4.17 |

**Supplementary Table 3.** Statistics for differentially methylated sites across assays.

|  |  |  |  |  |  |  |  |
| --- | --- | --- | --- | --- | --- | --- | --- |
| Number of Common Sites for all Assays | 2298846 | Assay |  |  |  |  |  |
| Common Sites with 5X for all Assays (C5X Sites) | 1928536 |  |  |  |  |  |  |
| DM Sites in 3 or more platforms on C5X sites (DM4+) | 29802 | EM-Seq | Methyl-Seq | Nanopore | SPLAT | TrueMethyl | TruSeq |
| Percentage of all common sites with 5X Coverage |  | 97% | 96% | 100% | 96% | 97% | 86% |
| Number of DM Sites for this assay on C5X Sites (DMA) |  | 74054 | 67621 | 26868 | 76591 | 59516 | 87170 |
| Percentage DMA unique to this platform |  | 26% | 26% | 17% | 29% | 22% | 42% |
| Percentage of DMA sites in DM4+ |  | 36% | 38% | 56% | 35% | 42% | 27% |
| Percentage of DM4+ in DMA sites |  | 90% | 86% | 51% | 89% | 85% | 79% |

**Supplementary Table 4.**
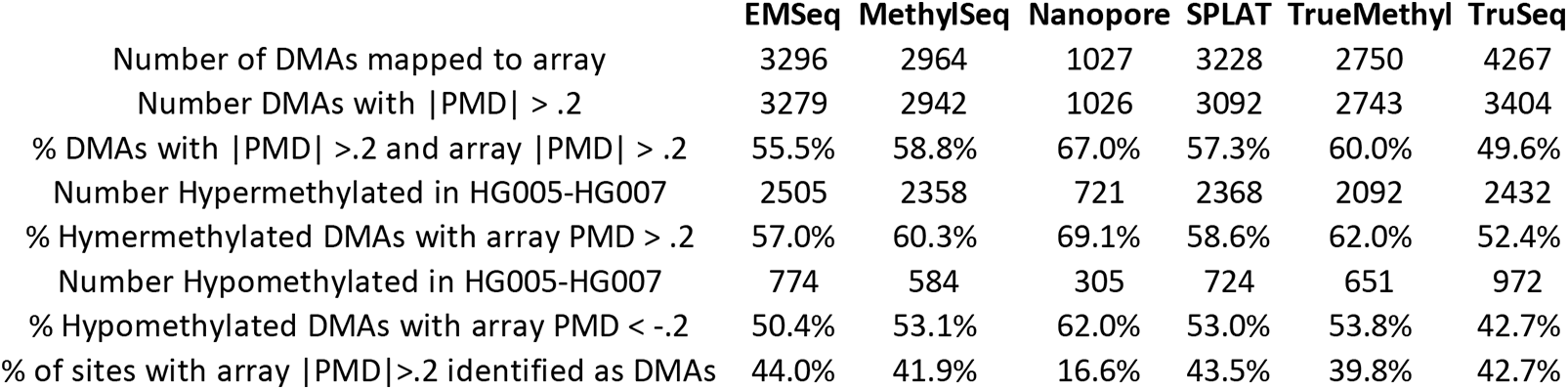
Concordance between assays of differentially methylated sites per assay (DMAs) with respect to microarray sites. PMD = Percent Methylation Difference, calculated as an absolute value.

**Supplementary Table 5.**
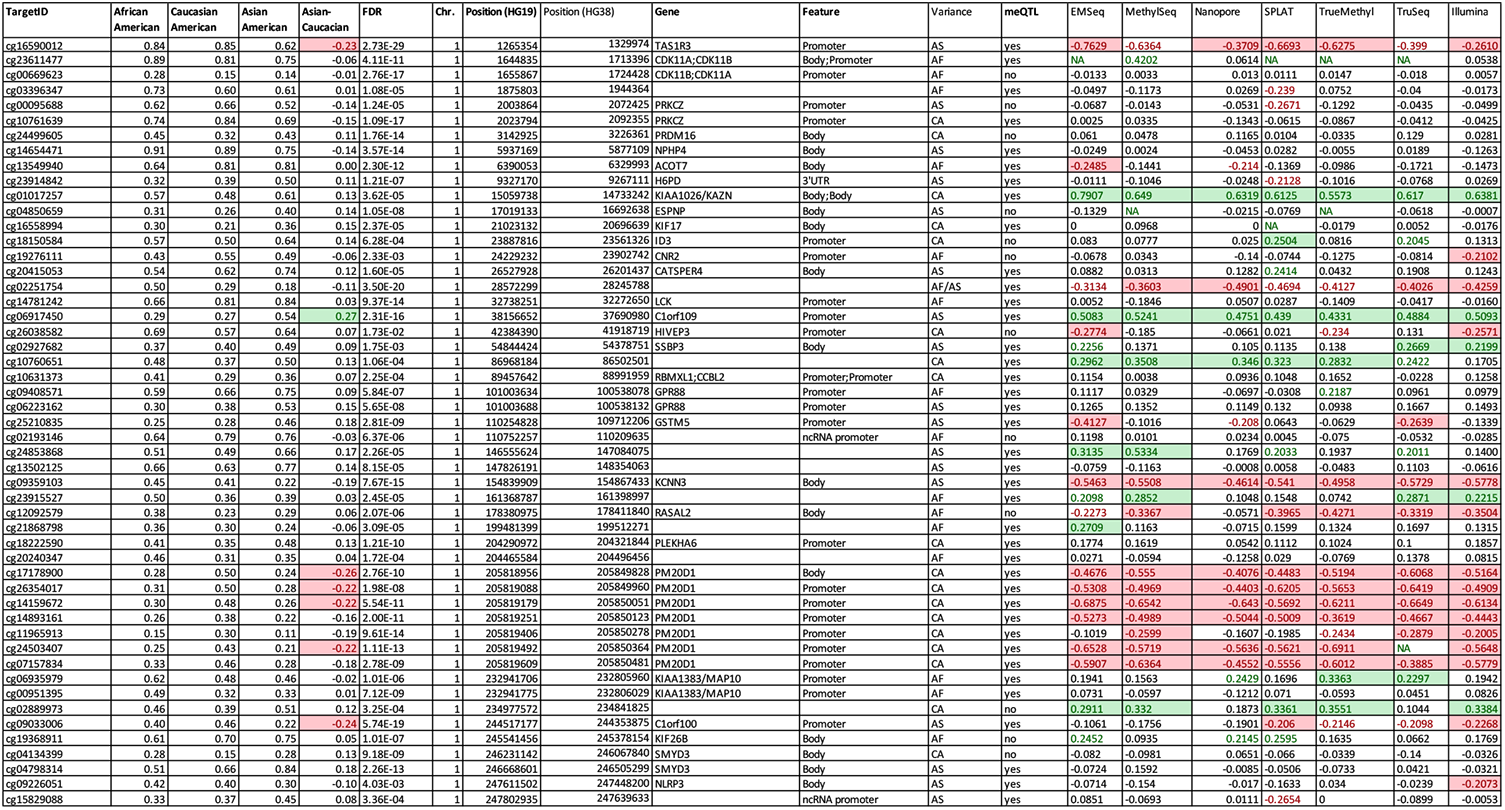
Population Variance agreement. A total of 52 CpGs on chromosome 1 that had been identified as differentially methylated between ethnic populations were annotated and compared for concordance of differential signal between microarray and sequencing data.

**Supplementary Table 6.** LC-MS/MS quantification of dC, dmC and dhmC in fmoles from digested genomic DNA (HG001-HG007) samples. For the detection of 5hmC a second higher volume injection was performed. The two dmC quantification values correspond to the two injections. Percentage of 5hmC in these samples is very low and below the limit of detection of the method.

| fmoles of different forms identified |  |  |  |  |  |  |  |  |  |
| --- | --- | --- | --- | --- | --- | --- | --- | --- | --- |
|  |  | Name | dC | dmC | dmC | dhmC | dmC/dC | dhmC/dmC | dhmC/dC |
| Biological Replicate 1 | HG001 | CP_01 | 5680.727 | 156.3333 | 2143.964 | 0.922163 | 2.75% | 0.04% | 0.0012% |
|  |  | CP_01 | 5721.555 | 157.2296 | 2212.077 | 0.966647 | 2.75% | 0.04% | 0.0012% |
|  | HG002 | CP_02 | 6134.437 | 206.1124 | 2915.523 | 1.078334 | 3.36% | 0.04% | 0.0012% |
|  |  | CP_02 | 6097.877 | 206.3662 | 2893.051 | 0.969016 | 3.38% | 0.03% | 0.0011% |
|  | HG003 | CP_03 | 6676.031 | 212.3023 | 2979.854 | 0.756934 | 3.18% | 0.03% | 0.0008% |
|  |  | CP_03 | 6742.651 | 211.5162 | 2948.914 | 0.739288 | 3.14% | 0.03% | 0.0008% |
|  | HG004 | CP_04 | 5223.132 | 188.2691 | 2588.429 | 0.592667 | 3.60% | 0.02% | 0.0008% |
|  |  | CP_04 | 5224.774 | 191.126 | 2560.265 | 0.56839 | 3.66% | 0.02% | 0.0008% |
|  | HG005 | CP_05 | 5487.878 | 192.3814 | 2680.448 | 0.828852 | 3.51% | 0.03% | 0.0011% |
|  |  | CP_05 | 5523.128 | 193.3392 | 2646.98 | 0.784192 | 3.50% | 0.03% | 0.0010% |
|  | HG006 | CP_06 | 5962.204 | 217.7679 | 2979.255 | 0.724408 | 3.65% | 0.02% | 0.0009% |
|  |  | CP_06 | 6041.553 | 217.7672 | 3002.681 | 0.686946 | 3.60% | 0.02% | 0.0008% |
|  | HG007 | CP_07 | 5819.142 | 205.9319 | 2860.277 | 0.884028 | 3.54% | 0.03% | 0.0011% |
|  |  | CP_07 | 5733.883 | 204.7114 | 2814.214 | 0.937631 | 3.57% | 0.03% | 0.0012% |
| Biological Replicate 2 | HG001 | CP_08 | 3620.176 | 99.23043 | 1362.303 | 0.646334 | 2.74% | 0.05% | 0.0013% |
|  |  | CP_08 | 3674.493 | 98.73453 | 1356.403 | 0.582917 | 2.69% | 0.04% | 0.0012% |
|  | HG002 | CP_09 | 2835.62 | 91.63515 | 1259.922 | 0.537728 | 3.23% | 0.04% | 0.0014% |
|  |  | CP_09 | 2872.229 | 92.00553 | 1250.31 | 0.520239 | 3.20% | 0.04% | 0.0013% |
|  | HG003 | CP_10 | 2832.307 | 85.18032 | 1167.688 | 0.370097 | 3.01% | 0.03% | 0.0010% |
|  |  | CP_10 | 2864.241 | 86.2764 | 1175.466 | 0.351737 | 3.01% | 0.03% | 0.0009% |
|  | HG004 | CP_11 | 2987.18 | 104.5185 | 1423.736 | 0.548236 | 3.50% | 0.04% | 0.0013% |
|  |  | CP_11 | 2989.671 | 104.0718 | 1431.635 | 0.452427 | 3.48% | 0.03% | 0.0011% |
|  | HG005 | CP_12 | 2999.949 | 102.7485 | 1410.854 | 0.489238 | 3.43% | 0.03% | 0.0012% |
|  |  | CP_12 | 3048.796 | 103.0123 | 1399.723 | 0.54343 | 3.38% | 0.04% | 0.0013% |
|  | HG006 | CP_13 | 3258.307 | 113.3027 | 1535.199 | 0.48512 | 3.48% | 0.03% | 0.0011% |
|  |  | CP_13 | 3199.5 | 111.8209 | 1519.138 | 0.476289 | 3.49% | 0.03% | 0.0011% |
|  | HG007 | CP_14 | 3324.166 | 113.9253 | 1545.66 | 0.634276 | 3.43% | 0.04% | 0.0014% |
|  |  | CP_14 | 3266.074 | 112.9173 | 1530.64 | 0.60005 | 3.46% | 0.04% | 0.0014% |
| Blank |  | CP_15 | 0 |  | 0.681001 |  |  |  |  |
|  |  | CP_15 | 0 |  | 0.689081 |  |  |  |  |

